# PATTY corrects open-chromatin bias for improved bulk and single-cell CUT&Tag profiling

**DOI:** 10.1101/2025.09.02.673784

**Authors:** Shengen Shawn Hu, Zhangli Su, Lin Liu, Qingying Chen, Megan C. Grieco, Mengxue Tian, Anindya Dutta, Chongzhi Zang

## Abstract

Precise profiling of epigenomes is essential for better understanding chromatin biology and gene regulation. Cleavage Under Targets & Tagmentation (CUT&Tag) is an efficient epigenomic profiling technique that can be performed on a low number of cells and at the single-cell level. With its growing adoption, CUT&Tag datasets spanning diverse biological systems are rapidly accumulating in the field. CUT&Tag assays use the hyperactive transposase Tn5 for DNA tagmentation. Tn5’s preference toward accessible chromatin alters CUT&Tag sequence read distributions in the genome and introduces open-chromatin bias that can confound downstream analysis, an issue more substantial in sparse single-cell data. We show that open-chromatin bias extensively exists in published CUT&Tag datasets, including those generated with recently optimized high-salt protocols. To address this challenge, we present PATTY (Propensity Analyzer for Tn5 Transposase Yielded bias), a comprehensive computational method that corrects open-chromatin bias in CUT&Tag data by leveraging accompanying ATAC-seq. By integrating transcriptomic and epigenomic data using machine learning and integrative modeling, we demonstrate that PATTY enables accurate and robust detection of occupancy sites for both active and repressive histone modifications, including H3K27ac, H3K27me3, and H3K9me3, with experimental validation. We further develop a single-cell CUT&Tag analysis framework built on PATTY and show improved cell clustering when using bias-corrected single-cell CUT&Tag data compared to using uncorrected data. Beyond CUT&Tag, PATTY sets a foundation for further development of bias correction methods for improving data analysis for all Tn5-based high-throughput assays.

## Introduction

Epigenetic mechanisms control gene expression and determine cell identity by altering chromatin states without changing DNA sequence^1^. Profiling epigenomic features, including transcription factor (TF) bindings and histone modifications (HM), across cell types, is important for understanding the molecular mechanisms of health and disease^2^. A series of high-throughput assays has been developed to generate epigenomic profiles. Advent in 2007^3,4^, ChIP-seq is the first and so far the most widely used epigenomic profiling technique using next-generation sequencing. ChIP-seq uses sonication or micrococcal nuclease (MNase) to digest chromatin, requiring a large number of cells and its performance depends on antibody quality and chromatin digestion efficiency. ChIP-exo^5^ can generate base-pair resolution TF binding patterns, but requires protein-DNA crosslinking, high-quality antibodies, and precise exonuclease control, limiting its usability. CUT&RUN (Cleavage Under Targets and Release Using Nuclease) uses protein A fused with MNase to target antibody and improves efficiency by only digesting chromatin associated with the target protein’s DNA binding sites^6^. CUT&Tag (Cleavage Under Targets and Tagmentation) was then developed to further improve the specificity and efficiency of epigenomic profiling^7^. CUT&Tag utilizes protein A (or G) fused with Tn5 transposase to specifically cleave and tag DNA at protein binding sites *in situ*, and can generate precise epigenomic profiles from native chromatin without cross-linking.

CUT&Tag has rapidly become popular because it is highly sensitive, often achieves strong signal-to-background, and has streamlined experimental protocols, making it particularly useful for samples with limited materials. Because of its high efficiency and *in situ* nature, CUT&Tag requires low DNA input, and can even be performed on a single-cell level^7^. The versatility of CUT&Tag has been further expanded through innovative modifications and improvements, leading to the development of novel applications, such as CUTAC^8^, TIP-seq^9^, and HiCuT^10^. Single-cell multi-omic joint profiling techniques were developed adapting modified CUT&Tag protocols to profile multiple histone modifications together with other modalities in the same cell, such as scCUT&Tag-pro^11^ for profiling histone modifications coupled with surface protein abundance, (Droplet) Paired-Tag^12,13^ for profiling histone modifications and transcriptomes in parallel, CUT&Tag2for1^14^, Multi-CUT&Tag^15^, and nano-CUT&Tag^16,17^ for profiling multiple histone modifications, RNA polymerase II, and chromatin accessibility. Spatial-CUT&Tag^18^ was also developed to spatially profile genome-wide histone modifications with a spatial resolution. Assays with CUT&Tag adoption are rapidly growing, with an increasing number of studies using it to investigate epigenetic mechanisms across species, biological systems, and diseases.

To accurately characterize epigenomic and chromatin states, computational analysis should be carefully designed to account for various noises and potential biases in the data generated by these high-throughput techniques. Tn5 transposase used in assays such as CUT&Tag can introduce two conceptually distinct types of biases: intrinsic cleavage bias and open-chromatin bias. Intrinsic cleavage bias arises from DNA sequence preferences of Tn5 transposase and manifests at the base-pair resolution^19^, whereas open-chromatin bias reflects the preferential activity of Tn5 transposase in accessible chromatin regions independent of DNA sequence^20–22^. While this accessibility preference constitutes the intended biological signal in ATAC-seq, it can represent a potentially confounding factor in CUT&Tag, where the goal is to profile antibody-targeted protein-DNA interactions such as histone modifications or transcription factor binding. As a result, open-chromatin bias in CUT&Tag can lead to spurious signal enrichment where the actual target mark is absent. Peak calling is commonly used for signal detection when analyzing CUT&Tag data. MACS2^23^ and SICER^24^ are widely used statistical model-based ChIP-seq peak calling tools, designed for narrow TF peaks and broad HM signals, respectively. SEACR^25^ was developed specifically to analyze CUT&RUN data. However, these conventional peak calling methods were not designed to correct the complicated biases caused by Tn5. We previously developed SELMA to characterize and correct Tn5’s intrinsic enzymatic cleavage bias in ATAC-seq^19^. SELMA primarily models DNA sequence-related bias and is effective for correcting fine-scale distortions in chromatin accessibility profiles including footprint patterns. However, SELMA is not designed to model biases driven by open chromatin, which is the intended signal in ATAC-seq but a potentially confounding factor in CUT&Tag. Bias correction for CUT&Tag is still challenging because the open-chromatin bias is dependent on chromatin context and cannot be adequately corrected by sequence-based models like SELMA alone. Additionally, for the highly sparse single-cell CUT&Tag data, the bias can be more substantial compared to bulk data. Innovative computational methods are thus urgently needed to accurately characterize and correct Tn5 biases in CUT&Tag data.

Here, we present PATTY (<u>P</u>ropensity <u>A</u>nalyzer for <u>T</u>n5 <u>T</u>ransposase <u>Y</u>ielded bias) to correct biases in CUT&Tag data at both bulk and single-cell levels, using ATAC-seq as a control. After observing widespread open-chromatin bias across published CUT&Tag datasets, we design a comprehensive strategy to benchmark multiple machine learning models and optimize the bias correction approach, and implement PATTY as a pre-trained factor-specific bias correction tool. We show that PATTY can correct open-chromatin bias from CUT&Tag data for both active (e.g., H3K27ac) and repressive (e.g., H3K27me3 and H3K9me3) histone modifications and generate more biologically meaningful results. We experimentally validated PATTY’s performance on H3K9me3 CUT&Tag. We further develop a computational framework built on PATTY to correct open-chromatin bias in single-cell CUT&Tag data, and improve single-cell clustering results.

## Results

### Tn5 open-chromatin bias affects the H3K27me3 CUT&Tag signal

The true histone modification profile in a given cell type/state should be invariant across different epigenomic assays. Therefore, we first compared the published genome-wide profiles of H3K27me3, a repressive histone modification, in the human K562 cell line, generated by CUT&Tag and ChIP-seq. While their peaks largely overlap, a substantial subset (18% on average) of CUT&Tag peaks is unique to CUT&Tag and does not overlap with ChIP-seq peaks (Figure 1a, b). We then examined these CUT&Tag-unique H3K27me3 signals. To our surprise, we observed a clear enrichment of H3K27me3 CUT&Tag signal near the transcription start sites (TSSs) of the actively transcribed genes (Figure 1c, d). As a repressive histone modification, H3K27me3 should not mark active gene promoters, where clear enrichments of nascent transcribed RNA and RNA Polymerase II (RNAPII) signals are observed (Figure S1a-c), confirming active transcription. In contrast, this H3K27me3 pattern is not present in ChIP-seq, consistent with the transcriptionally active state of these regions (Figure 1e). This widespread H3K27me3 signal observed across active transcribed promoter regions indicates a potential artifact in CUT&Tag. Interestingly, the ATAC-seq signal for chromatin accessibility showed a similar pattern to the CUT&Tag signal at these regions (Figure 1f). Indeed, the H3K27me3 peaks that uniquely appear in CUT&Tag but not in ChIP-seq are enriched at active gene promoter regions and ATAC-seq determined open chromatin regions (Figure S1d, e). Considering that active gene promoters are open chromatin and that both ATAC-seq and CUT&Tag use Tn5 to access chromatin and cleave DNA, we argue that the CUT&Tag signals for H3K27me3 at active promoter regions are false signals associated with open-chromatin bias caused by Tn5 in the assay.

**Figure 1.**
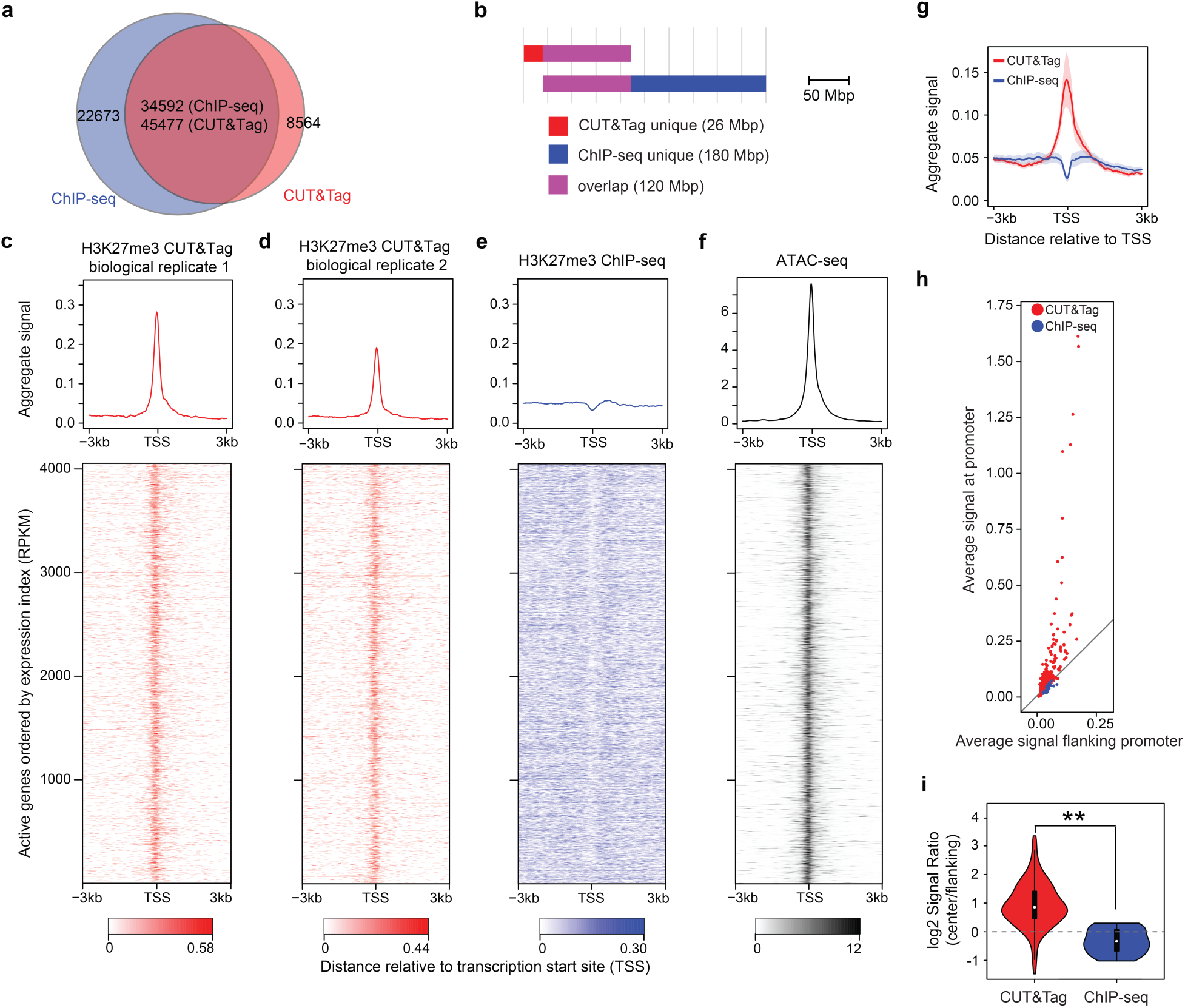
Aberrant CUT&Tag signals for H3K27me3. (**a**) Overlapping peak counts between CUT&Tag and ChIP-seq for H3K27me3 in human K562 cell line. Peaks were called separately from ChIP-seq and from CUT&Tag. The number of ChIP-seq peaks overlapping CUT&Tag peaks and the number of CUT&Tag peaks overlapping ChIP-seq peaks were reported separately, as they differ due to different peak boundaries. (**b**) Genomic lengths of overlapping and non-overlapping peak regions between CUT&Tag and ChIP-seq for H3K27me3 in K562. (**c**-**f**) Signal patterns across active promoter regions in K562 cells, for two biological replicates of H3K27me3 CUT&Tag (**c**,**d**), H3K27me3 ChIP-seq (**e**), and ATAC-seq (**f**). The upper panels are the normalized aggregate signal patterns. The lower panels are heatmaps of the signal patterns at promoter regions (TSS ±3kb) of the actively transcribed genes. Rows correspond across heatmaps. (**g**) Aggregate H3K27me3 signal patterns at the promoter regions of 1000 ubiquitously expressed genes of 277 human CUT&Tag datasets (red) and 30 ENCODE human ChIP-seq datasets (blue). The shades represent the 95% confidence interval. (**h**) Average H3K27me3 signal levels at the proximal promoter region (TSS ± 300bp, y-axis) against the flanking regions (300bp – 3kb from TSS on both sides, x-axis) of the ubiquitously expressed genes for CUT&Tag (red) and ChIP-seq (blue) datasets. Each data point represents a dataset. (**i**) Distributions of log2-scaled ratio between the proximal promoter (center) region and the flanking region average signals for CUT&Tag and ChIP-seq datasets. **, p < 0.01, by one-sided Wilcoxon signed-rank test.

We then surveyed more H3K27me3 CUT&Tag datasets to assess how commonly the open-chromatin bias occurs, especially with the recent protocol improvements to remove or reduce open-chromatin bias by using high-salt washes. We further collected 277 H3K27me3 CUT&Tag datasets across various human cell types published since 2024, most of which reported using updated high-salt wash protocols, and examined the signals at the promoter regions of a group of ubiquitously expressed genes, where H3K27me3 is not expected to be present. We observed strong enrichment of H3K27me3 CUT&Tag signals near the TSSs, a pattern clearly distinct from ENCODE^26^ ChIP-seq data (Figure 1g). To quantitatively assess this promoter enrichment pattern across datasets, we compared the average normalized signal in the proximal promoter regions (TSS ±300 bp) of the ubiquitously expressed genes to that in the flanking regions (300 bp–3 kb on both sides of the TSS) of these genes for each dataset. Most CUT&Tag datasets exhibited substantially higher signal levels at proximal promoters relative to flanking regions, whereas ChIP-seq datasets showed minimal differences (Figure 1h). We further quantified this pattern using the proximal/flanking signal ratio as a metric, and found a significant difference between CUT&Tag and ChIP-seq datasets (Figure 1i). These observations suggest that open-chromatin bias is a widespread issue across H3K27me3 CUT&Tag data, regardless of experimental protocol improvements.

### Defining a ground truth benchmark for true and false H3K27me3 marked regions

To further characterize the H3K27me3 CUT&Tag signals and to create a ground truth benchmark that can be used for model training, we curated a set of true and false H3K27me3 CUT&Tag signal patterns using collected CUT&Tag data, incorporating prior biological knowledge. We assumed that true H3K27me3 should be associated with non-expressed genes, and as reciprocal conjugate modifications, H3K27me3 and H3K27ac should not coexist at the same genomic locus in any autosome in a homogeneous cell population. Based on these assumptions, we defined a true H3K27me3-marked region as a reproducible H3K27me3 CUT&Tag peak covered region around non-expressed genes, with no H3K27ac CUT&Tag sequence reads. In the meantime, we defined a false H3K27me3-marked region as a reproducible H3K27me3 CUT&Tag peak covered region around highly expressed genes while overlapping with H3K27ac CUT&Tag peaks (Figure 2a, b, Methods). We collected 17 high-quality H3K27me3 CUT&Tag datasets and generated such defined sets of true and false H3K27me3-marked regions in the human K562 cell line. At the 200bp nucleosomal resolution, we obtained 1428 true (285.6kb) and 1231 false (246.2kb) H3K27me3 CUT&Tag signal-marked regions. As expected, we found that the false H3K27me3 CUT&Tag regions exhibited significantly higher ATAC-seq signal levels than the true H3K27me3-marked regions, suggesting biases due to open chromatin (Figure 2c).

**Figure 2.**
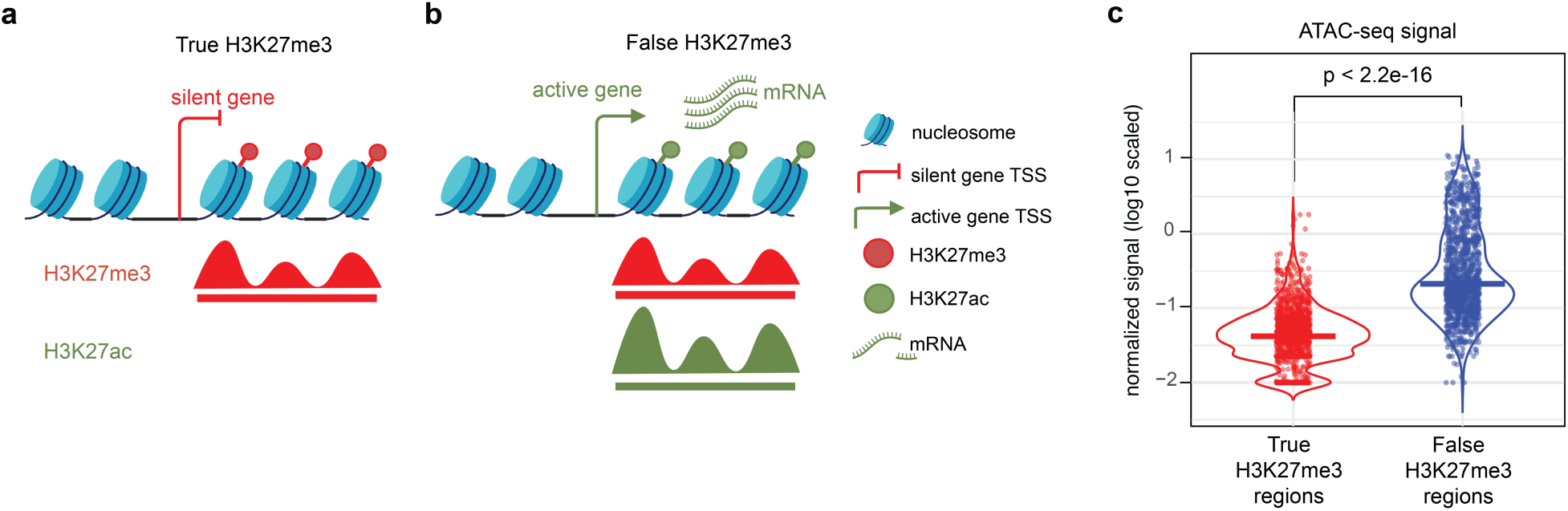
Curation of true and false H3K27me3 CUT&Tag signals. (**a**, **b**) Schematics of genomic regions with true (**a**) and false (**b**) H3K27me3 CUT&Tag signals. Red and green represent repressive and active chromatin features, respectively. (**c**) Distribution of chromatin accessibility level (ATAC-seq signal) on the true (red) and false (blue) H3K27me3 CUT&Tag signal regions. Each data point represents a 200-bp region. The centerline represents the median value. The P-value is calculated by one-sided Wilcoxon rank sum test.

### Evaluating machine-learning models to predict true H3K27me3-marked regions with multi-modal features

To identify true H3K27me3 marked regions from CUT&Tag data considering the open-chromatin bias, we sought to develop a supervised machine-learning model to classify true and false H3K27me3 signal regions from multi-modal features on the pre-defined ground truth benchmark data from human K562 cells (Figure 3a, Methods). For each 200bp region in the genome, we considered the following features as candidate covariates: the 10bp-resolution normalized signal patterns across the 1kb window centered at the 200bp bin for H3K27me3 CUT&Tag (the primary signal), ATAC-seq (to represent an empirical readout of open-chromatin context), and IgG (to represent a negative control^8^), as well as the 200bp one-hot encoded DNA sequence (to represent intrinsic sequence preference at base-pair resolution^27,28^).

**Figure 3.**
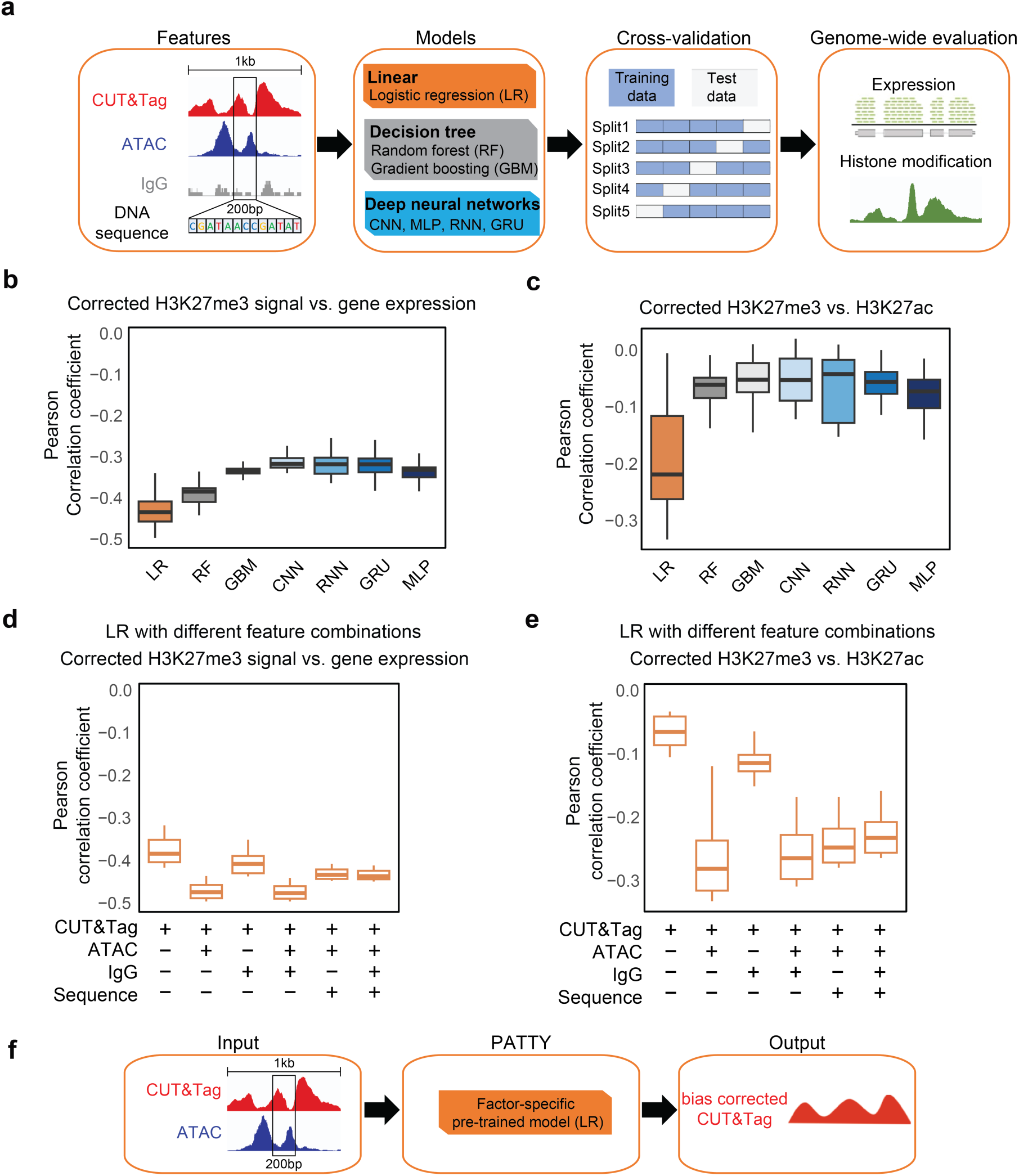
Machine learning models to classify true and false H3K27me3 regions with multi-dimensional features. (**a**) Workflow of model construction and evaluation. (**b**) Pearson correlation between the model derived H3K27me3 CUT&Tag score and gene expression across all gene promoter bins in the genome. (**c**) Pearson correlation between the model derived H3K27me3 CUT&Tag score and H3K27ac ChIP-seq signal level across all bins with CUT&Tag reads in the genome. Each data point in a boxplot represents a CUT&Tag sample from the model with a particular feature combination. (**d**, **e**) Similar to (**b**, **c**) respectively, for only the logistic regression model with various feature combinations labeled below the plots. The centerline, bounds of the box, vertical line bottom, and vertical line top of the boxplots represent the median, 25th to 75th percentile range, 25th percentile − 1.5 × interquartile range (IQR), and 75th percentile + 1.5 × IQR, respectively. Results from different feature combinations were combined into the same box. (**f**) Schematic of PATTY workflow.

We employed several commonly used machine-learning models for this task, including panelized Logistic Regression (LR), Random Forest (RF), and Generalized Boosted Model (GBM), as well as state-of-the-arts deep neural network models, including Convolutional Neural Networks (CNN)^29–31^, Multilayer Perceptron (MLP), Recurrent Neural Network (RNN), and Gated Recurrent Unit (GRU), to identify the best-performing model (Figure S2a-c, Methods). We first assessed the models’ performance using a 5-fold cross-validation on the pre-defined true/false signal regions. Using all feature combinations, we found that the performance varies across different models and different feature combinations, and deep learning-based models seem to have better performance than other models (Figure S2d).

Next, we trained each model using the whole benchmark set of true/false H3K27me3 regions, and applied the trained model to the whole genome to determine whether any genomic region with H3K27me3 CUT&Tag signal has a true or false H3K27me3 mark in human K562 cells. We used an orthogonal, biology-informed approach to evaluate the model performance: We calculated the genome-wide correlation between the model prediction score (true/false H3K27me3 signal) and either RNA-seq or H3K27ac ChIP-seq signal level, and assess the model performance based on the biological ground truth that as a repressive histone modification, H3K27me3 should be negatively correlated with both gene expression and H3K27ac signal across the genome in the same cell type. As a result, the logistic regression (LR) model showed consistently the best performance, i.e., yielding the highest negative correlation between the model-predicted H3K27me3 and both gene expression (Figure 3b, S2e) and H3K27ac signals (Figure 3c, S2f). Moreover, among all feature combinations, including only the CUT&Tag and ATAC-seq signals yielded a better performance of the LR model than all other feature combinations as represented by the highest negative correlation with both gene expression (Figure 3d) and H3K27ac signal (Figure 3e). Neither IgG nor DNA sequence can further improve the model performance, while ATAC-seq is essential to the true H3K27me3 determination, indicating the importance of considering open-chromatin bias in obtaining the correct H3K27me3-marked regions (Figure 3d, e). These results suggested that the LR model using CUT&Tag and ATAC-seq signal features is an optimal approach to correct open-chromatin bias and generate accurate genome-wide H3K27me3-marked regions for H3K27me3 CUT&Tag data analysis. Therefore, we implement this model as a computational tool called Propensity Analyzer for Tn5 Transposase Yielded bias (PATTY), for improved analysis of histone modification CUT&Tag data by correcting open-chromatin bias (Figure 3f).

### PATTY corrects open-chromatin bias and improves H3K27me3 and H3K27ac CUT&Tag analysis

We applied PATTY to generate the corrected H3K27me3 profiles for the two H3K27me3 CUT&Tag datasets samples shown in Figure 1c,d, and found that the false signal pattern at the active gene promoter regions caused by the open-chromatin bias was corrected throughout the genome (Figure 4a-f), exemplified at two active gene loci, NUP214^32^ (Figure 4g) and GCSH^33^ (Figure 4h). We further showed that the PATTY-corrected H3K27me3 CUT&Tag profile is more negatively correlated with gene expression (Figure 4i) and H3K27ac profile (Figure 4j), compared to the H3K27me3 profile generated by peak-calling with macs2, with or without considering IgG control, but without bias correction. Meanwhile, the H3K27me3 signal patterns near the repressed gene promoter regions were not much affected after PATTY correction (Figure 4g, h, S3). As a reference, the effect of PATTY correction on H3K27me3 CUT&Tag signals was also demonstrated across genes associated with our curated true and false H3K27me3-marked regions (Figure S4).

**Figure 4.**
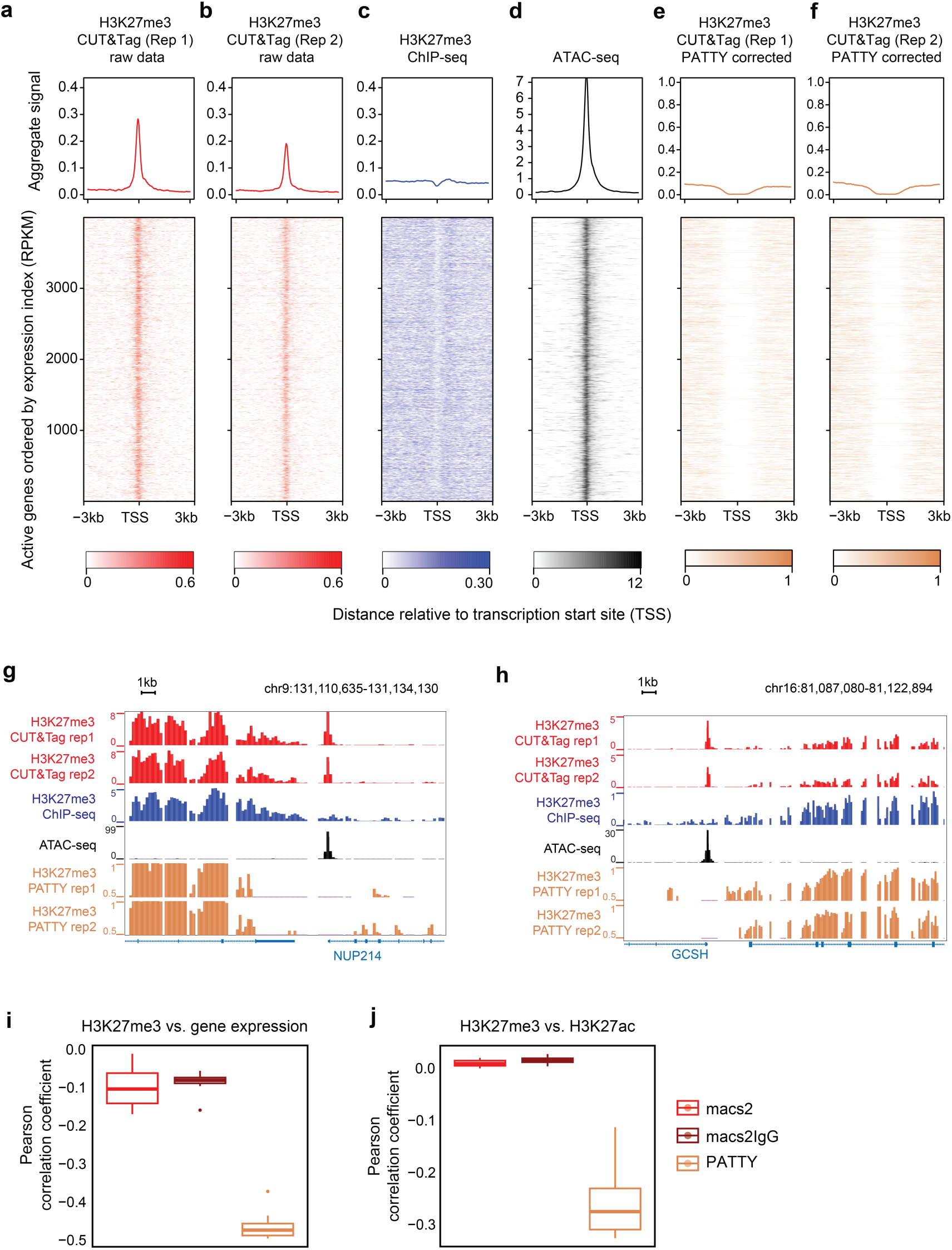
PATTY correction of open-chromatin bias in H3K27me3 CUT&Tag data. (**a**-**f**) H3K27me3 CUT&Tag signal patterns across active gene promoter regions before (**a**, **b**) and after (**e**, **f**) bias correction by PATTY. ChIP-seq (**c**) and ATAC-seq (**d**) signals across the same regions are shown for reference (similar to Figure 1e**,f**). (**g**, **h**) Genome browser snapshots at the loci of two active genes, NUP214 (**g**) and GCSH (**h**), in K562. (**i**, **j**) Pearson correlation between H3K27me3 CUT&Tag signal, processed by various methods, and gene expression across all gene promoter bins in the genome (**i**) or H3K27ac ChIP-seq signal across all bins with CUT&Tag reads in the genome (**j**), in the K562 cell line. Each data point in a boxplot represents a different H3K27me3 CUT&Tag sample.

It has been reported that modified CUT&Tag experimental protocols, such as using high-salt wash steps, can help reduce open-chromatin bias in H3K27me3 CUT&Tag data^20^. We then specifically tested the performance of PATTY on H3K27me3 data generated from such modified CUT&Tag protocols, including high-salt CUT&Tag^20^ (Figure 5a, b) and CUTAC^8^ (Figure 5c, d). Interestingly, although the open-chromatin bias was supposed to be reduced by these modified protocols, the H3K27me3 profiles after PATTY correction still attained higher negative correlations with both gene expression (Figure 5a, c) and H3K27ac (Figure 5b, d), indicating that there are still residue biases in the data even with modified experimental protocols, confirming what we observed (Figure 1g-i), and PATTY is still able to correct the bias computationally and further improve the data interpretations. These results suggest that PATTY can successfully correct open-chromatin bias in H3K27me3 CUT&Tag data and improve signal detection.

**Figure 5.**
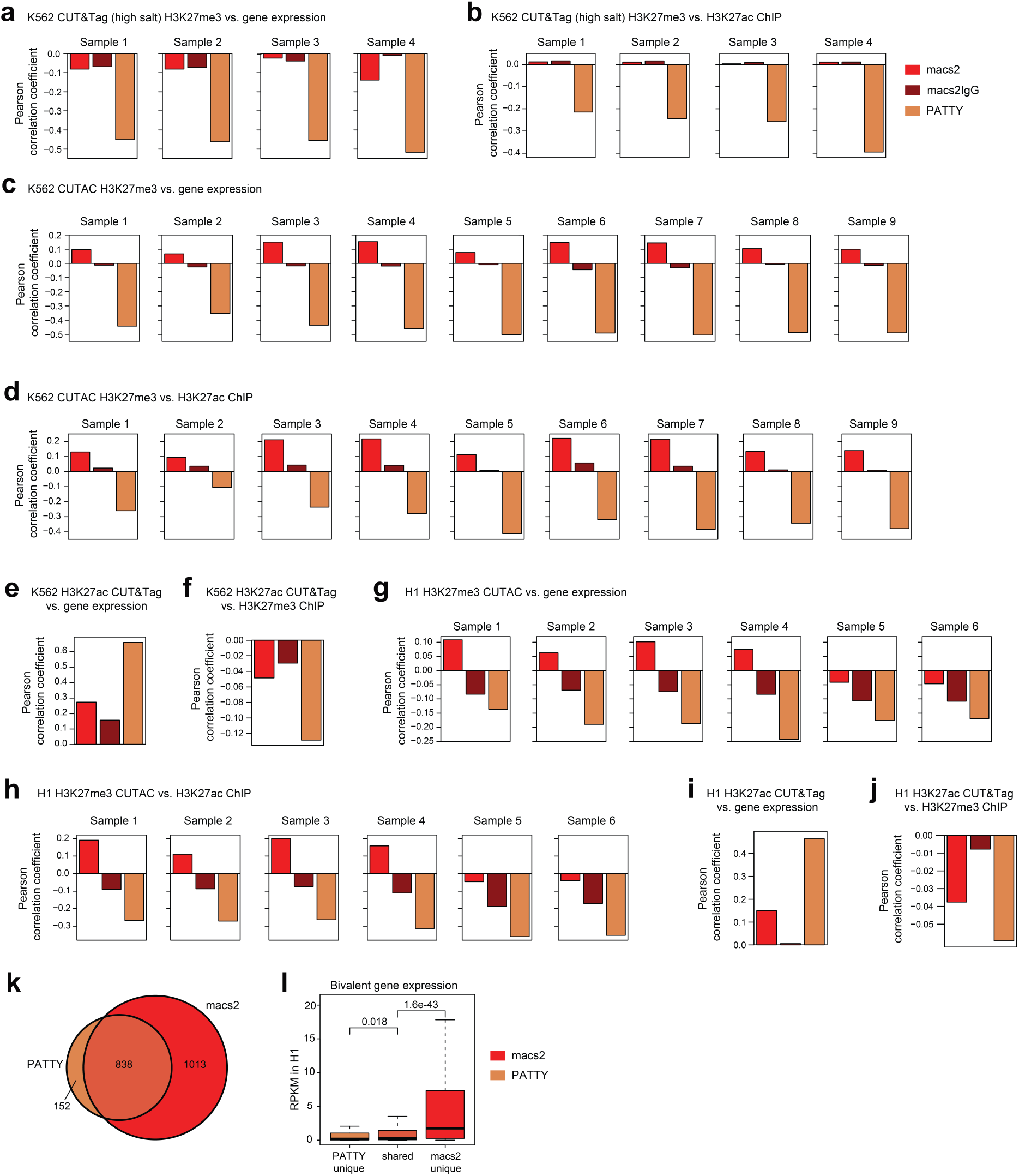
PATTY correction of CUT&Tag biases for H3K27me3 and H3K27ac across various samples and cell types. (**a**-**d**) Pearson correlation between processed H3K27me3 CUT&Tag signal and gene expression across all genes (**a**, **c**) or H3K27ac ChIP-seq signal across all 200-bp bins in the genome (**b**, **d**) in the K562 cell line. CUT&Tag samples in (**a**, **b**) were prepared in high-salt concentration. Samples in (**c**, **d**) were generated using the CUTAC assay. (**e**, **f**) Pearson correlation between processed H3K27ac CUT&Tag signal and gene expression across all genes (**e**) or H3K27me3 ChIP-seq signal across all 200-bp bins genome-wide (**f**), in K562. (**g**-**j**) Similar to (**a**, **b**, **e**, **f**) but for H1 human embryonic stem cell (hESC) line. (**k**) Bivalent genes identified in H1 hESC using PATTY-corrected data and using uncorrected macs2 peaks. (**l**) Expression levels in H1 hESC for three sets of bivalent genes in (**k**). P-values were calculated by one-sided Wilcoxon rank-sum test.

We then tested PATTY’s ability to correct open-chromatin bias in CUT&Tag data for H3K27ac, an active histone modification. Unlike repressive chromatin marks, which generally do not co-occur with open chromatin, the Tn5-induced signal may confound the true active histone modification signal at the same locus, making bias correction more challenging than for repressive marks. Because the machine learning framework in PATTY is agnostic to whether the CUT&Tag is for active or repressive marks, we applied the same workflow for H3K27ac. Specifically, we constructed the benchmark by defining the true H3K27ac-marked regions as H3K27ac CUT&Tag peak regions around highly expressed genes with no H3K27me3 CUT&Tag reads, and false H3K27ac-marked regions as H3K27ac CUT&Tag peak-covered regions around non-expressed genes while overlapping with H3K27me3 CUT&Tag peaks. In this way, we curated a set of true and false H3K27ac CUT&Tag signal regions using collected CUT&Tag data in K562, trained the H3K27ac PATTY model using the H3K27ac CUT&Tag and ATAC-seq signal patterns as input features, and applied the trained model to generate the genome-wide corrected H3K27ac CUT&Tag profile, with the correction effect across genes associated with the curated true and false H3K27ac regions shown in Figure S5. As a result, we observed that the PATTY-corrected H3K27ac profile has an increased positive correlation with gene expression (Figure 5e), and a higher negative correlation with its reciprocal conjugate histone modification, H3K27me3 (measured by ChIP-seq for orthogonality), compared to conventional macs2 peak calling without bias correction (Figure 5f). These results indicate that PATTY can correct open-chromatin biases existing in CUT&Tag for both active and repressive histone marks.

### Pretrained PATTY model can correct CUT&Tag biases across different cell types

Next, we examined whether the pre-trained PATTY model using our pre-defined ground truth datasets from one cell type can correct the open-chromatin bias in CUT&Tag for the same histone modification in other cell types and improve data analysis universally. We applied the PATTY model trained with data in K562 to several CUT&Tag datasets generated in the human embryonic stem cell line H1, and observed consistently improved analysis results compared with conventional peaking calling without bias correction, for both H3K27me3 (Figure 5g, h) and H3K27ac (Figure 5i, j), as demonstrated using the same metrics such as correlation with gene expression (Figure 5g, i) or with the reciprocal conjugate histone modification (Figure 5h, j). These data suggest that using the model pre-trained from one cell type, PATTY is powerful for correcting open-chromatin bias and improving analysis of CUT&Tag data from different cell types. This result also indicates that the machine-learning model in PATTY captures the CUT&Tag intrinsic open-chromatin bias specific to each histone modification rather than any cell-type-specific information.

H3K4me3/H3K27me3 bivalent chromatin is a unique epigenetic feature in stem cells and pluripotent progenitor cells. We then assessed the effect of PATTY correction in the identification of bivalent chromatin domains and associated genes in H1. We defined a gene as bivalent gene if its promoter region overlapped with both H3K4me3 and H3K27me3 signals. We compared the sets of bivalent genes identified using the uncorrected H3K27me3 signal with those obtained using the PATTY-corrected H3K27me3 signal. Although PATTY correction resulted in fewer bivalent genes identified genome-wide (990 vs. 1851; Figure 5k), many of those 1013 “bivalent” genes detected only with the uncorrected signal (macs2) are unlikely to represent true bivalent loci, as they exhibited high expression levels (Figure 5l), contradicting to the canonical view that bivalent genes are poised and either silent or transcribed at a very low level^34^. In contract, the bivalent genes uniquely identified after PATTY correction exhibited minimal to low transcription levels, similar to those bivalent genes consistently identified with or without PATTY correction (Figure 5l), consistent with the expected behavior of canonical bivalent genes. These results suggest that PATTY can improve the identification of biologically meaningful bivalent domains and associated genes in relevant cell types.

### PATTY corrects bias and improves CUT&Tag data analysis for H3K9me3

After demonstrating PATTY’s performance with extensive public data for H3K27me3 and H3K27ac, we next sought to evaluate PATTY’s effectiveness on our own experimental data for H3K9me3, a histone modification associated with inactive heterochromatin. We used a similar strategy to curate a ground-truth benchmark dataset of true/false H3K9me3-marked regions in K562 using CUT&Tag data for reciprocal conjugate histone modifications H3K9me3 and K3K9ac. Specifically, we defined the true H3K9me3-marked regions as reproducible H3K9me3 CUT&Tag peak regions around non-expressed genes with no H3K9ac CUT&Tag reads, and false H3K9me3-marked regions as reproducible H3K9me3 CUT&Tag peak-covered regions around highly expressed genes while overlapping with H3K9ac CUT&Tag peaks. We used this benchmark dataset to train a PATTY model for H3K9me3, and observed the effect of PATTY correction across genes associated with our curated true and false H3K9me3-marked regions (Figure S6). We performed a CUT&Tag experiment for H3K9me3 in the HCT116 cell line and tested the performance of PATTY on this in-house dataset. Similar to H3K27me3, the open-chromatin bias caused a clearly false enrichment of H3K9me3 CUT&Tag signal at active gene promoter regions (Figure 6a), a pattern absent in ChIP-seq (Figure 6b). As expected, the false signals caused by open-chromatin bias (Figure 6c) were corrected by PATTY (Figure 6d), exemplified by several gene loci (Figure 6e-g). Meanwhile, H3K9me3 CUT&Tag signals at non-expressed gene loci were less affected by the open-chromatin bias, and therefore, they were similar to ChIP-seq signals and not altered by PATTY correction (Figure 6e-g, S7). We also examined PATTY’s performance using the same genome-wide correlation metrics and found that the PATTY-corrected H3K9me3 CUT&Tag profile is negatively correlated with both gene expression (Figure 6h) and the profile of H3K9ac (Figure 6i). Such negative correlations were not detected if the H3K9me3 CUT&Tag data only underwent MACS2 peak calling without bias correction (Figure 6h, i). Therefore, our in-house experiment for H3K9me3 validated the effectiveness of PATTY in correcting open-chromatin bias in CUT&Tag data.

**Figure 6.**
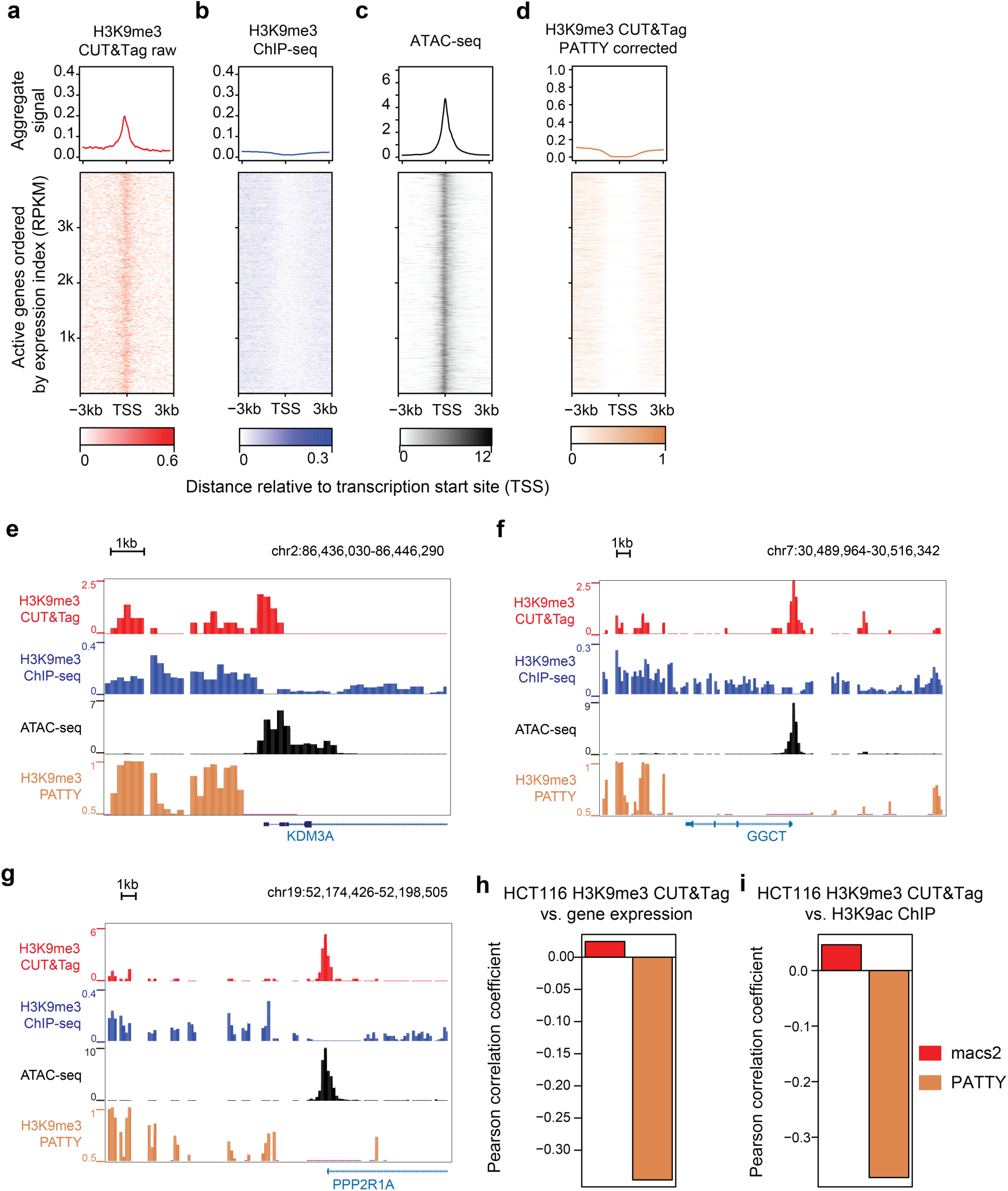
PATTY correction of open-chromatin bias in CUT&Tag for H3K9me3. (**a**-**d**) H3K9me3 CUT&Tag signal patterns across active gene promoter regions before (**a**) and after (**d**) bias correction by PATTY. ChIP-seq (**b**) and ATAC-seq (**c**) signals across the same regions are shown for reference (similar to Figure 4a-f), in human HCT116 cell line. (**e**-**g**) Genome browser snapshots at the loci of several active genes in HCT116. (**h**, **i**) Pearson correlation between H3K9me3 CUT&Tag signals and gene expression across all gene promoter bins in the genome (**i**) or H3K27ac ChIP-seq signal across all bins with CUT&Tag reads in the genome (**j**), in the HCT116 cell line.

### PATTY improves cell-type clustering on single-cell CUT&Tag data

Recent innovations in single-cell CUT&Tag enable the detection of epigenomic profiles in each of hundreds or thousands of cells^11,12,14–17,35,36^. Single-cell CUT&Tag data are often highly sparse, and most histone modification events in individual cells can only be represented by a single DNA fragment, which could amplify the effect of the Tn5-caused open-chromatin bias. We collected H3K27me3 and H3K27ac single-cell CUT&Tag datasets from three independent studies that employed different single-cell multiomic co-assays, nano-CT^17^, scCUT&Tag-pro^11^, and Paired-Tag^12^, each of which not only profiled multiple histone marks but also reported cell-type annotation for each individual cell to serve as the ground truth. Specifically, cell-type annotations were derived from cell surface markers for scCUT&Tag-pro^11^ and from integrated multi-modality information for nano-CT^17^ and Paired-Tag^12^, respectively. Using two groups of cells annotated as CD8+ T cells and monocytes from the scCUT&Tag-pro dataset^11^ as an example, we observed enrichments of H3K27me3 CUT&Tag signals at the promoter regions of ubiquitously expressed genes, where H3K27me3 should not occur (Figure S8a, b), and differentially expressed genes between the two cell types (Figure S8c-f), where H3K27me3 should not occur in the respective cell type highly expressing these genes (Figure S8d, e), suggesting open-chromatin bias affecting this single-cell dataset from exhibiting specific H3K27me3 signals to differentiate cell types.

We then assessed whether PATTY could correct the bias and improve the analysis of these single-cell CUT&Tag datasets. To reduce data sparsity and for PATTY to work genome-wide seamlessly, we employed a meta-cell approach, to represent the signal profile of each individual cell with the average signal profile of that cell and its 10 nearest neighboring cells in the top 50 principal component (PC) space. We then applied PATTY to each cell’s meta-cell-smoothed CUT&Tag signal using ATAC-seq from the same cell system, and generated a PATTY-corrected single-cell CUT&Tag data matrix. We then performed unsupervised cell clustering using both the PATTY-corrected and the uncorrected single-cell CUT&Tag data matrices in parallel. We evaluated clustering accuracy by comparing the results from PATTY-corrected and uncorrected data against the ground-truth cell-type annotations. Using the H3K27me3 nano-CT dataset as an example, and visualizing all cells under the same UMAP projection derived from the ground truth annotation reported in the original publication, we colored the cells by their cluster assignments based on the uncorrected data (Figure 7a) or the PATTY-corrected data (Figure 7b). We found that clustering based on the PATTY-corrected data showed better agreement with the ground-truth cell-type annotations (Figure 7c), as evidenced by more distinct color separations on different cell groups on the UMAP visualization. A similar trend was observed for the H3K27ac nano-CT dataset (Figure 7d-f). We then quantified this observation using the adjusted Rand index (ARI; higher values indicate better agreement between 2 clustering results on the same dataset) by comparing each clustering result with the ground-truth cell-type assignments. Indeed, clustering based on the PATTY-corrected data yielded higher ARI values than clustering based on the uncorrected data, for both H3K27me3 (Figure 7g) and H3K27ac (Figure 7h). Similarly, we observed consistent improvements in clustering performance after PATTY correction on the scCUT&Tag-pro (Figure 7i, j, S9a-f) and Paired-Tag (Figure 7k, l, S9g-l) datasets, for both H3K27me3 (Figure 7i, k, S9a-c, g-i) and H3K27ac (Figure 7j, l, S9d-f, j-l) as well. These data suggest that PATTY correction on single-cell CUT&Tag data can improve cell clustering accuracy.

**Figure 7.**
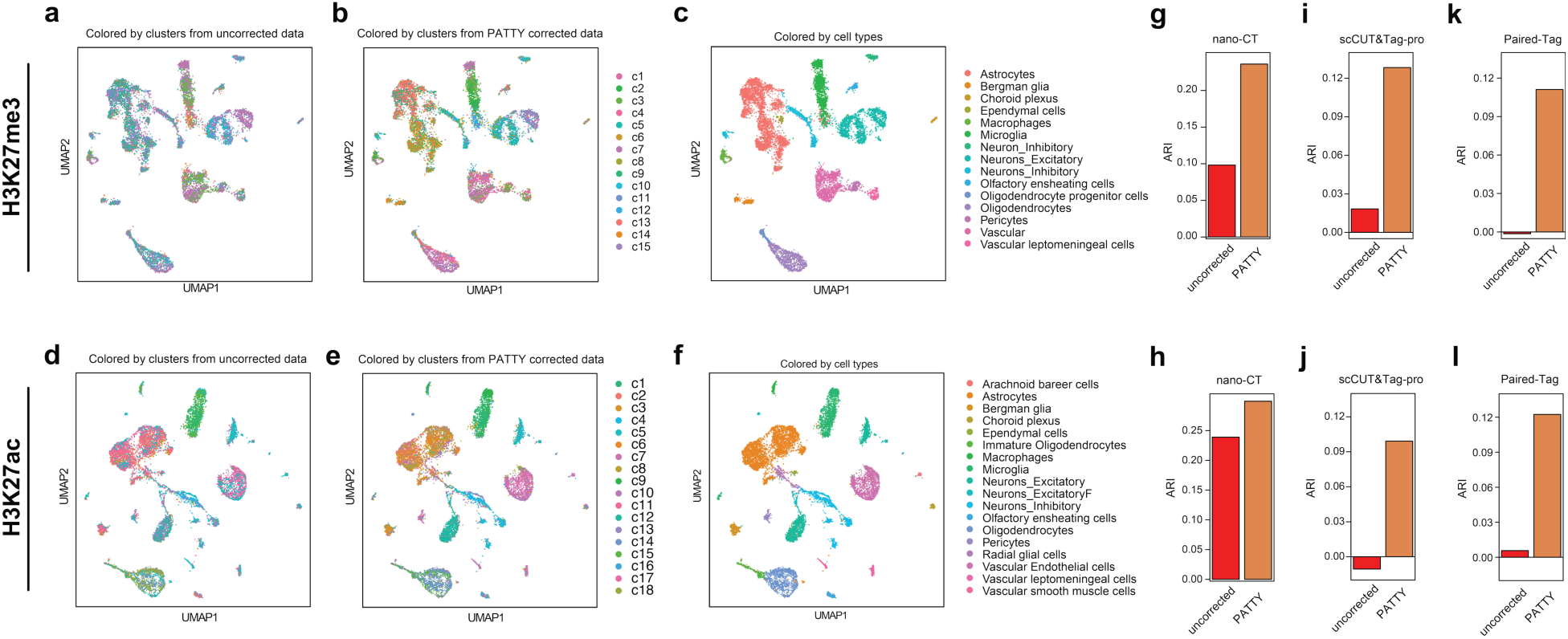
Improved cell clustering on single-cell CUT&Tag after PATTY correction. (**a**-**f**) Visualization of H3K27me3 (**a**-**c**) and H3K27ac (**d**-**f**) single-cell nano-CT data using UMAP projection reported in the original publication. Cells are colored by cluster assignments based on uncorrected data (**a**, **d**), cluster assignments based on PATTY-corrected data (**b**, **e**), or the published cell-type annotations (ground truth) (**c**, **f**). (**g**-**l**) Adjusted Rand index (ARI) between the ground-truth cell-type annotation and the clustering results from uncorrected and PATTY-corrected data, for nano-CT (**g**, **h**), scCUT&Tag-pro (**i**, **j**), and Paired-Tag (**k**, **l**) datasets for H3K27me3 (**g**, **i**, **k**) and H3K27ac (**h**, **j**, **l**), respectively.

Next, we further examined the effects of PATTY correction on single-cell CUT&Tag data. In the scCUT&Tag-pro dataset, PATTY-corrected H3K27me3 signals exhibited the expected differential patterns at the differentially expressed genes between monocytes and CD8+ T cells (Figure S8g, h). For the Paired-Tag dataset, which includes a gene expression modality, we quantified the correlation between CUT&Tag signals and nearby gene expression across the genome for each annotated cell type. Consistent with biological expectations, PATTY-corrected H3K27me3 signals exhibited stronger negative correlations with gene expression, whereas PATTY-corrected H3K27ac signals showed stronger positive correlations with gene expression, across all annotated cell types (Figure S10a, b). Since nano-CT enables joint profiling of H3K27me3 and H3K27ac in the same cells, we additionally evaluated whether PATTY correction can improve multimodal integration. We performed weighted nearest neighbor (WNN) analysis by integrating the two modalities, and found that WNN-based clustering using PATTY-corrected data showed improved ARI with the ground-truth cell-type annotations compared to using uncorrected data (Figure S10c). Together, these data further support the effectiveness of PATTY for bias correction in single-cell CUT&Tag data and the benefit of PATTY generalizes across single-cell CUT&Tag platforms and experimental designs.

## Discussion

As CUT&Tag has been increasingly used for epigenomic profiling, characterization and correction of biases in CUT&Tag data have become inevitable. We have shown the widespread existence and severe impact of open-chromatin bias in CUT&Tag, including both bulk and single-cell datasets generated with the latest protocols. We demonstrated the ability of PATTY in correcting open-chromatin bias for both bulk and single-cell histone modification CUT&Tag data, using ATAC-seq as control, and validated the results using both public and in-house experimental data. PATTY has been implemented as an open-source software tool pre-trained for H3K27me3, H3K27ac, and H3K9me3 CUT&Tag. It can be used for any cell type, biological systems, either bulk or single cell, without re-training.

We attribute the observed bias to Tn5 transposase, which plays a critical role in the CUT&Tag assay, especially in generating fragment libraries by inserting sequencing adapters into the DNA, i.e., tagmentation. Since Tn5 transposase prefers chromatin-accessible regions^37^, utilizing Tn5 in CUT&Tag inevitably results in some read enrichment towards open-chromatin regions, regardless of the feature of the target protein of CUT&Tag experiments^20^. This bias is intrinsic to Tn5 and therefore cannot be completely eliminated by experimental protocol optimization, as demonstrated in recently published datasets. We use ATAC-seq to control this bias, because ATAC-seq provides a direct measurement of chromatin accessibility, which reflects the propensity of Tn5 to access and tagment DNA due to chromatin openness. In the PATTY framework, ATAC-seq is not used as a conventional background control for peaking calling (such as the chromatin input control for ChIP-seq), but rather as an empirical readout of the open-chromatin-derived component that confounds CUT&Tag signals, because the quantitative effects caused by Tn5 transposase in CUT&Tag and ATAC-seq are not as simple as a signal-control relationship, and more sophisticated modeling is required. Therefore, we incorporate ATAC-seq signal as input covariates in the machine-learning models to estimate and remove the open-chromatin-associated contribution to CUT&Tag signal patterns, while preserving the antibody-targeted signal of interest. Importantly, this is achieved through modeling local signal patterns, rather than simple background subtraction or signal-noise statistical modeling, as we believe this approach allows for more accurate disentanglement of chromatin accessibility-driven and target-specific signals. Indeed, we have shown that PATTY outperforms state-of-the-art peak-calling methods like macs2.

A key step in developing a well-performing machine learning model is to prepare appropriate, high-quality training data. In this work, it boils down to determining the ground truth epigenomic pattern that we can use for model training. In the PATTY framework, the models are specific to each factor/histone modification due to the specific biological features of each epigenomic mark. While the training data are from one common cell line, once the model is trained, it can be used in any cell type for that same epigenomic mark. In this work, we curated ground truth benchmark datasets for H3K27me3, H3K27ac, and H3K9me3 using publicly available data based on orthogonal biological knowledge. For other widely studied histone marks, such as H3K4me3, H3K4me1, and H3K36me3, however, curating appropriate ground truth benchmark datasets may not be as straightforward using the same strategy, because they do not have a well-established reciprocal conjugate mark with opposing functions for identifying mutually exclusive chromatin domains. Moreover, histone marks like H3K4me3 and H3K4me1 can be associated with both actively transcribed and silent/poised genes, making gene expression information unsuitable for reliable ground-truth curation. Interestingly, when we applied the pretrained PATTY model for H3K27ac (as a proxy for active chromatin marks) directly to H3K4me3, H3K4me1, and H3K36me3 CUT&Tag data without training a new model or fine tuning, we still observed a consistent increased correlation between histone modification signals and nearby gene expression after PATTY correction (Figure S11). While this may suggest that the pretrained H3K27ac PATTY model might capture some general open-chromatin bias and could potentially be applicable to other active histone marks, we remain cautious and do not recommend this practice without further rigorous validation. When a rigorously defined ground truth dataset is available for a histone mark, we can apply the same framework to train a PATTY model for bias correction for that mark.

The model design for PATTY highlights the importance of biological knowledge in computational biology method development. When orthogonal biological information is used for model evaluation, popular black-box deep learning models do not perform as well as simple, more interpretable logistic regression. This is consistent with a recent study showing that deep learning models did not outperform linear models in single-cell gene perturbation prediction^38^. Therefore, choosing an appropriate model is crucial. The model selection should always consider biological or scientific information for the specific goal, and should not solely rely on blindly increasing model complexities. When comparing different feature combinations, it is also interesting that considering DNA sequence features does not necessarily yield a better performance. This is possibly because the sequence bias information has already been implicitly included in the control ATAC-seq data^19^. When evaluating model performance, we use orthogonal and biologically meaningful metrics such as correlation with gene expression and with reciprocal conjugate histone modifications, rather than relying on correlation or similarity with ChIP-seq data. This is because we consider CUT&Tag as a more advanced technique, or at least an alternative, to ChIP-seq. Therefore, we view the two techniques as parallel and do not treat ChIP-seq as a ground truth or benchmark against with CUT&Tag should be judged.

While we curated the ground truth benchmark data under the assumption that H3K27ac and H3K27me3 rarely co-occur at the same genomic locus due to their antagonistic mechanism^39^, we note that apparent signal overlap between these two histone marks can also reflect cellular heterogeneity in bulk datasets, where the signals of the two marks originate from distinct cell populations rather than true co-occupancy within the same cells. This scenario can be particularly relevant during differentiation or in progenitor-like populations where cell states are transient and heterogeneous. In our study, model training and most evaluations were performed in established cell lines (K562, H1, and HCT116), which are generally less heterogeneous than primary tissues or differentiating cell populations and therefore less susceptible to strong population-mixing effects. In addition, the ground truth datasets were curated by integrating dozens of datasets and selecting the most reproducible regions, further minimizing the potential confounding effects of cellular heterogeneity. Nevertheless, one should always remain cautious when interpreting signals derived from heterogeneous bulk cell populations.

Tn5 transposase is a powerful enzyme in epigenetics research. In addition to ATAC-seq and CUT&Tag, Tn5 transposase has also been used in other high-throughput assays for whole genome sequencing (LIANTI^40^), chromatin interaction detection (HiChIP^41^ and HiCAR^42^), and spatial epigenomic profiling (epigenomic MERFISH^43^), etc. Similar to CUT&Tag, the Tn5 preference to open chromatin could induce potential biases in data from those assays as well. Modeling and correcting open-chromatin bias and other potential enzymatic biases in these high-throughput data remain a significant task in computational biology. Our work sets a foundation for open-chromatin bias correction in other data types, and more work can be built to help improve the analysis for meaningful biological interpretation.

## Methods

### High-throughput sequencing data collection and processing

CUT&Tag, ChIP-seq, ATAC-seq, and IgG data were processed as follows: Raw sequencing reads were aligned to the GRCh38 (hg38) reference genome with bowtie2^44^ (v2.2.9) (-X 2000 for paired-end data). Low-quality reads (MAPQ < 30) were discarded. For paired-end sequencing data, reads with two ends aligned to different chromosomes (chimeric reads) were discarded. For paired-end data, reads with identical 5’ end positions for both ends were regarded as redundant reads and discarded. Reads mapped to mitochondrial DNA (mtDNA reads) were discarded. To generate the genome-wide signal track for downstream analysis (i.e., scan signal pattern on genome-wide bins, and generate signal heatmap across gene promoters), we extended reads from their 5’end to 146 base pairs (bp), piled them up, and normalized the piled-up signal by total read count in the sample (i.e., RPM). For CUT&Tag and ChIP-seq data, SICER (v2)^24^ peaks were detected with default parameters (used in Figure 1a, b, and curation of True/False benchmark regions). For H3K27ac CUT&Tag, H3K4me1 CUT&Tag, H3K4me3 CUT&Tag, H3K27ac ChIP-seq, and ATAC-seq data, macs2^23^ peaks were detected with additional parameters “-q 0.01 --nomodel --extsize 146”. For H3K27me3, H3K9me3, and H3K36me3 CUT&Tag and ChIP-seq data, macs2 peaks were detected with additional parameters “-q 0.01 –broad --nomodel --extsize 146”.

RNA-seq was processed as follows: Raw sequencing reads were aligned to the GRCh38 (hg38) reference genome with Hisat2 (v2.2.1)^45^. Low-quality reads (MAPQ < 30) were discarded. We then estimated the expression index (RPKM) using Stringtie (v2.1.5)^46^.

Exon-array data were processed with the “jetta” package^47^. We collected all exon array samples from the study (GSE19090) as the input data and used the “jetta.do.expression” function to estimate and normalize the expression index of samples. The average expression index from 3 replicates in the K562 cell line was used as the K562 gene expression.

To examine the performance of recent CUT&Tag data with the latest protocols, we collected all human H3K27me3 CUT&Tag datasets from GEO generated since 2024, in total 277 samples. For comparison, we also collected 30 H3K27me3 ChIP-seq datasets from the ENCODE project^26^ (generated from two different institutes). We selected a set of ubiquitously expressed genes as the top 1000 genes with the highest minimum expression index across all cell types (normalized expression from Exon array data) and examined the pattern and enrichment of all 277 samples on the promoter (TSS ±3kb) of these genes. For each H3K27me3 dataset, we calculated the average signal pattern across the promoter regions of all ubiquitously expressed genes. Then the average patterns from the 277 samples were summarized to generate Figure 1g. To quantify the level of signal enrichment in promoter regions, we calculated the average normalized signal level at promoter center region, defined as TSS ± 300bp, and flanking promoter region, defined as 300bp-3kb from TSS on both sides, respectively, for the 1000 ubiquitously expressed genes. The center and flanking scores for all 277 datasets were visualized as a scatter plot in Figure 1h. The log2 ratio of center and flanking scores was further compared (Figure 1i).

For single-cell RNA-seq data in human PBMC^48^, we downloaded the processed data and used Seurat^49^ (v5.3.0) for the data processing and analysis. The cell type information was inherited from published data. The PBMC-ubiquitously expressed genes (used in Figure S8a, b) were defined with sufficient average expression index in all PBMC cell types (normalized average expression > 0.2). Two cell types with sufficient cell numbers (CD8+ T cells and monocytes) were collected, and we performed differential expression analysis (fold change >=2 and adjusted p-value < 0.05) to capture differentially expressed genes between the two cell types used in Figure S8.

For single-cell CUT&Tag data, processed data (aligned fragments and Seurat/Signac R objects) were downloaded from the original studies^11,12,17^. We used the ArchR package (v1.0.1)^50^ for single-cell data processing. All high-quality cells provided were used. The top 25k tilling bins with the highest cross-cell signal variance were collected and split into 200bp bins for the downstream analyses (referred to as high-var bins below).

### Curation of true and false H3K27me3/H3K27ac marked regions

H3K27me3 and H3K27ac CUT&Tag datasets in the K562 cell line were collected from Reference^7^, the study with the most datasets, without data from other publications to avoid potential batch effects. Peak calling analysis was performed on the collected CUT&Tag datasets with SICER (v2). Peaks were then split into 200bp tiling bins as candidate H3K27me3 or H3K27ac regions.

Considered as the ground truth, the true H3K27me3 marked regions were selected based on the following criteria: 1) H3K27me3 marked regions should not be actively transcribed. So we selected candidate H3K27me3 regions (in 200bp bins obtained from SICER peaks) from each CUT&Tag H3K27me3 dataset that are overlapped with the promoter (± 1kb from transcriptional start site, TSS) or gene body regions of non-expressed genes, defined as those with 0 RPKM detected from K562 RNA-seq data (8,032 genes in total). 2) H3K27me3 and H3K27ac should be mutually exclusive in a homogeneous cell population. So we filtered out the candidate H3K27me3 regions that have any H3K27ac CUT&Tag reads from any H3K27ac CUT&Tag dataset. 3) H3K27me3 regions should be reproducible from many CUT&Tag samples. So we selected those remaining candidate H3K27me3 regions (200bp bins) that reoccurred in at least 14 out of the total 17 H3K27me3 CUT&Tag samples as the final true H3K27me3 regions.

Accordingly, the false H3K27me3 regions were selected based on the following criteria: 1) H3K27me3 signals detected in actively transcribed regions are false. So we selected candidate H3K27me3 regions (in 200bp bins obtained from SICER peaks) from each CUT&Tag H3K27me3 dataset that overlapped with the promoter or gene body regions of the highly expressed genes, defined as the top 8,032 genes (on par with the non-expressed genes for selecting the true H3K27me3 regions) ranked by the expression level (in RPKM) in K562. 2) H3K27me3 and H3K27ac should be mutually exclusive in a homogeneous cell population. Thus, we further selected those candidate H3K27me3 regions that overlapped with H3K27ac CUT&Tag peaks as the false H3K27me3 regions.

The true H3K27ac regions were selected as the H3K27ac SICER peak regions that overlapped with the promoter regions of the highly expressed genes, and had no H3K27me3 CUT&Tag read detected in the candidate region or the ± 500bp flanking regions from any H3K27me3 CUT&Tag dataset. The false H3K27ac regions were selected as the H3K27ac SICER peak regions that overlapped with the promoter regions of the non-expressed genes, and also overlapped with H3K27me3 CUT&Tag SICER peaks from any H3K27me3 CUT&Tag sample.

The true H3K9me3 regions in K562 were selected as the H3K9me3 SICER peak regions that 1) have greater than or equal to 5 reads in both K562 H3K9me3 CUT&Tag samples, 2) do not overlap with the gene body or ± 10kb flanking regions of the highly expressed genes, and 3) have no H3K9ac CUT&Tag reads detected in the regions. The false H3K9me3 regions were selected as the H3K9me3 SICER peak regions from any H3K9me3 CUT&Tag sample that overlapped with the gene body or ± 10kb flanking regions of the highly expressed genes, and had at least 5 reads from any H3K9ac CUT&Tag sample.

In order to show the signal patterns analysis for curated ground-truth benchmark regions (Figures S4-S6), we first collected the curated true and false 200-bp regions. We then defined “true-region-related genes” as genes whose TSS ±3 kb window has at least 1 bp overlap with any true region, and “false-region-related genes” as genes whose TSS ±3 kb window has at least 1 bp overlap with any false region. For each gene set, we aggregated signals in a ±3 kb window centered at the TSS to generate the aggregate profile and heatmap visualizations.

We only included histone modification signals on the 22 autosomes and assumed homozygous chromatin states. The X chromosome was excluded to avoid potential confounding effects arising from X chromosome inactivation. The overlapping status between the two region sets was determined by BEDtools^51^ with at least 1bp overlap.

### Classifying true and false regions with different machine learning models and various genomic feature combinations

We built supervised models to predict the true/false classification (1/0) for the histone modification marked regions, employing a collection of machine learning techniques, including panelized logistic regression (LR), random forest (RF), gradient boosting machine (GBM), and deep neural networks (dNN)., and using various combinations of genomic features, including CUT&Tag signal pattern, ATAC-seq signal pattern, IgG signal pattern, and one-hot encoded DNA sequence (Figure 3a). For each candidate region (a 200bp bin) as an observed sample, the 10bp-resolution signal pattern (normalized DNA fragment pile-up count) across a 1kb region centered on the bin center was considered as a feature vector, for CUT&Tag, ATAC-seq, and IgG. During model training, signal patterns from different H3K27me3 CUT&Tag datasets on the same candidate region were considered independent observed samples. When multiple signal patterns from CUT&Tag, ATAC-seq, or IgG profiles were selected as features in the dNN models, the multiple signal patterns were aligned by genomic locations as a matrix feature. The one-hot encoded DNA sequence for the entire 200bp bin was considered as the sequence feature. In the dNN models, if signal pattern features (i.e., CUT&Tag, ATAC-seq, or IgG) and DNA sequence features were both included in the model, the signal features and sequence features were considered as two branches, and their flattened outputs were concatenated later.

The panelized logistic regression (LR) model used elastic net penalty (alpha = 0.5), with maximum iterations set as 10,000. The decision tree number (n_estimators) of the random forest (RF) model was set as 100. In LR, RF, and GBM models, all selected features were flattened and horizontally concatenated as the model input.

We tested four dNN architectures in the model exploration steps (Figure 3a, S2), including Convolutional Neural Networks (CNN), Multilayer Perceptron (MLP), Recurrent Neural Networks (RNN), and Gated Recurrent Unit (GRU). For all four models, an input layer with shape (200, 4) was defined for the one-hot encoded sequence features, where 200 represents the nucleotide composition in each bp across the 200bp bin. An input layer for the signal feature had a shape (100, N), where 100 represents the 100 data points for the 10bp-resolution signal across the 1kb region, and N, chosen from 1-3, represents the selected number of signal pattern features, e.g., N=1 for CUT&Tag only, N=2 for CUT&Tag + ATAC or CUT&Tag + IgG, and N=3 for CUT&Tag + ATAC + IgG.

For CNN (Figure S2a), 1) a 1D convolutional layer with 32 filters and ReLU activation was applied. L2 regularization was used to prevent overfitting. 2) A MaxPooling1D layer with a pool of size 2 was added to reduce the dimensionality and computational complexity of the feature maps, and to extract the most salient features. All these layers were applied to both the sequence and the signal branches. 3) The flattened outputs from both branches were concatenated. 4) A dense layer with 128 units and ReLU activation was added, and L2 regularization was applied again in this step. 5) A dropout layer for mitigating overfitting and a final dense layer with a single unit and sigmoid activation for binary classification (true/false labels) were added. The kernel size and dropout rate were tuned for optimized model performance.

For MLP (Figure S2b), 1) the input layers for both branches were passed through the flatten layer to convert the input tensors into one-dimensional vectors. 2) The vectors were then concatenated into a single vector. 3) The combined vector was fed into a dense layer with 128 units and ReLU activation. L2 regularization was applied to this layer to prevent overfitting. 4) The output from the first dense layer was then passed into a second dense layer with 64 units and ReLU activation, also with L2 regularization. 5) The final output layer was a dense layer with a single unit and a sigmoid activation function. The regularization coefficient was tuned for optimized model performance.

For RNN and GRU (Figure S2c), 1) the input layers were processed through an LSTM/GRU layer with 32 units and L2 regularization. The LSTM/GRU layer would capture temporal dependencies within both the sequence and signal branches. 2) The outputs from the two LSTM/GRU layers from the two branches were concatenated into a single vector, which was the input to a dense layer with a single unit, and a sigmoid activation function was applied to the concatenated output.

All dNN models were trained using Adam with a learning rate tuned for optimized performance evaluated by cross-validation (described below). The loss function was set to binary cross-entropy, which is suitable for binary classification tasks. We designed a 5-fold cross-validation to evaluate the generalization performance using accuracy metrics (Figure S2a). The parameters, including kernel size and dropout rate in CNN, as well as regularization coefficient and learning rate, were also tuned in this step. For each dNN model, the hyperparameter with the optimized performance was used for the genome-wide analyses (Figure 3b-e).

All dNN models were implemented with TensorFlow and Keras (v2.11.0) packages, and other models were implemented with the sklearn (v1.0.2) packages.

### Genome-wide evaluation of machine learning models

After training a model using the curated true/false datasets with cross-validation, we applied the trained model to the genome-wide 200bp bins and obtained a model prediction score for each bin (referred to as the corrected signal in Figure 3b-e). The model prediction score ranges from 0 to 1. A prediction score closer to 1 means the bin is more likely to have a true signal, and a prediction score closer to 0 means the bin is more likely not to have a true signal. We next collected all bins located within ± 1kb from any gene’s TSS (promoter bins) and examined the expression level of that gene whose promoter each bin is located in. We calculated the Pearson correlation coefficient between the corrected signal (model prediction score) and the associated gene expression across all promoter bins (e.g., Figure 3b, d). We assumed that repressive histone marks (e.g., H3K27me3, H3K9me3) should be negatively correlated with gene expression, and active histone marks (e.g., H3K27ac) should be positively correlated with gene expression. Therefore, we used the calculated correlation coefficient as a metric to evaluate the model’s performance. Because the RNA-seq data in the K562 cell line had been used for selecting true/false CUT&Tag signal regions as training data, here we used the exon array data in K562 as the gene expression measurement for this genome-wide evaluation, to maintain the orthogonality and rigor.

When calculating the model output’s correlation with other histone mark profiles (e.g., Figure 3c, e), we used the ChIP-seq signal for the other histone mark in the same cell line. For H3K27ac and H3K9me3 CUT&Tag data, bins with at least one read were included for the correlation calculation. For H3K27me3 CUT&Tag data, bins with at least one read in at least 6 samples were included for the calculation. For the macs2 results presented for comparison, a bin overlapped with a macs2 peak was assigned a score as the Macs2 generated fold enrichment value of that peak. The other bins that are not associated with any Macs2 peak were assigned a score of 0.

We further evaluated whether the pretrained H3K27ac PATTY model can be applied to other active histone modifications. Specifically, we applied the pretrained H3K27ac PATTY model to CUT&Tag datasets of H3K4me3, H3K4me1, and H3K36me3 using the same feature construction. We collected all bins located within ± 1kb from any gene’s TSS (promoter bins) and examined the expression level of that gene whose promoter each bin is located in. Similar to the H3K27ac evaluation strategy, we calculated the Pearson correlation coefficient between the corrected signal (PATTY score) and the associated gene expression across all promoter bins. We expect that each of the 3 histone marks (H3K4me3, H3K4me1, and H3K36me3) should be positively correlated with gene expression (Figure S11).

### Bivalent domain analysis

To evaluate bivalency in H1 ESC, we defined bivalent bins as genomic regions jointly marked by H3K4me3 and H3K27me3. For the uncorrected data, we called peaks from raw CUT&Tag using MACS2 with narrowPeak for H3K4me3 (--nomodel --extsize 146 -q 0.01) and broadPeak for H3K27me3 (--broad --nomodel --extsize 146 -q 0.01). For the PATTY-corrected data, we used the same MACS2-called H3K4me3 peaks, but defined H3K27me3 peaks as PATTY peaks using a stringent cutoff (PATTY score ≥ 0.9). We then defined bivalent bins as the genomic intersection between the H3K4me3 peak set and the H3K27me3 peak set. Finally, we defined bivalent genes as genes whose promoter region (TSS ±1 kb) overlaps any bivalent bins. Bivalent gene sets derived from uncorrected versus PATTY-corrected data were compared by categorization into PATTY-unique, shared, and MACS2-unique groups (Fig. 5k), and gene expression in H1 ESC (RPKM) was compared across groups using a two-sided unpaired Wilcoxon rank-sum test (Fig. 5l).

### PATTY method

Based on the model performance results, LR models were employed in the PATTY method. In practice, we implemented PATTY using the pre-trained LR model for each specific histone mark. The input of PATTY includes the mapped reads files (in BED format) of a CUT&Tag dataset and an ATAC-seq dataset from the same cell sample. PATTY will generate the signal tracks (for genome browser visualization) and run the LR model prediction across the genome to generate a 200bp resolution corrected signal profile. Any prediction score below 0.5 can be reduced to 0 for visualization purposes. Peak calling can be further performed on the PATTY-corrected signal profile.

### Signal patterns on active/silent gene promoter regions

The top 4,000 genes with the highest expression level (measured by RPKM value from RNA-seq) and having ATAC-seq signal in their promoter regions (TSS ± 3kb) in K562 cells were selected as K562 active genes. Genes with 0 RPKM in K562 RNA-seq data and with at least 5 reads in each dataset of both H3K27me3 CUT&Tag replicates and H3K27me3 ChIP-seq were selected as silent genes. For the raw CUT&Tag, ChIP-seq, and ATAC-seq signals, sequence depth-normalized signal (described in the data collection and processing section) was used for the aggregate plots, heatmaps, and genome browser visualizations. The PATTY corrected signal was used for the PATTY corrected plots.

### Single-cell PATTY score and evaluation

To evaluate PATTY’s performance on single-cell CUT&Tag data, we first developed a meta-cell approach to reduce the single-cell data sparsity before applying PATTY. In detail, we first calculated the pairwise Euclidean distances of all cell pairs in the top 50 high-variance principal component (PC) space from the ArchR preprocessed output. For each individual cell, the top 10 nearest cells (with the shortest distance) were selected as the neighboring cells. The scCUT&Tag signal patterns of these 10 neighboring cells and the target cell itself (11 cells in total) were merged together to represent the CUT&Tag signal pattern of the target cell. We collected publicly available (sc)ATAC-seq from the same cell types as the scCUT&Tag data for the ATAC-seq signal pattern, i.e., 10x Genomics PBMC scATAC-seq for scCUT&Tag-pro, ENCODE mouse frontal cortex ATAC-seq for Paired-Tag^26^, and the scATAC-seq component in the nano-CT experiments for the nano-CT scCUT&Tag data. To reduce the computing complexity, we treated the scATAC-seq data as bulk ATAC-seq data. Then, for each individual cell, we used the meta-cell adjusted CUT&Tag signal pattern and the bulk ATAC-seq signal pattern as the input for PATTY to correct bias and generate the signal of the cell on all candidate bins. Finally, we can generate a PATTY score matrix for all cells across all candidate bins as the bias-corrected scCUT&Tag data matrix.

To compare the raw data matrix and PATTY-corrected data matrix, we first conducted a PCA dimensional reduction using the top 10k high-variance bins. Next, we ran a k-means clustering on the PCA output, where k was set to the same number of cell types in the ground truth, which referred to the “L2” level cell type annotation in scCUT&Tag-pro^11^ and nano-CT^17^, and “dna_Anno_transfer” cell type in Paired-Tag^12^. Then, the clustering result was compared to the ground truth annotation, and an adjusted Rand index (ARI) was calculated to assess the clustering accuracy (Figure 7g-l, Figure S10c). We applied this ARI approach to both the uncorrected raw data matrix and the PATTY-corrected data matrix for a fair comparison. We further visualized the cell type annotation and clustering labels (from both uncorrected data and PATTY-corrected data) on the UMAP visualization.

For the nano-CT dataset, we performed weighted nearest neighbor (WNN) integration of H3K27me3 and H3K27ac measured in the same cells with shared barcodes (Figure S10c). Briefly, we first computed principal components (PCs) separately for each modality and retained the top 30 PCs per cell for each of H3K27me3 and H3K27ac (using either the uncorrected data matrix or the PATTY-corrected data matrix). We then restricted to cells present in both modalities (by barcode intersection). The two modality-specific PC embeddings were then collected and the WNN modality weights were learned using Seurat FindMultiModalNeighbors function. A joint embedding was constructed as a weighted combination of the two modality-specific PC embeddings. We then performed clustering on the joint embedding with the same methods for uni-modal data.

### Cell lines and culture

HCT116 p53-/- cell (referred to as wild-type, RRID: CVCL_S744) was a generous gift from Fred Bunz (Johns Hopkins)^52^. All cell lines were maintained in McCoy’s 5A-modified medium (Corning, no. 10-050-CV) supplemented with 10% fetal bovine serum. Cell lines were authenticated by STR method and routinely tested for mycoplasma contamination by PCR.

### H3K9me3 CUT&Tag library preparation

H3K9me3 CUT&Tag libraries were prepared using Active Motif CUT&Tag-IT Assay Kit (cat #2200) with H3K9me3 antibody (Diagenode #C15410193) following the manufacturer’s instructions. Specifically, 500,000 cells and 1 μg antibody were used per reaction.

## Data availability

The H3K9me3 CUT&Tag data in HCT116 is available at the Gene Expression Omnibus (GEO) with accession number GSE298565.

## Code availability

The source code of PATTY and scripts for all analyses of this work are available at Github: https://github.com/zang-lab/PATTY

## Acknowledgements

The authors thank Dr. Ye Zheng for helpful suggestions and critical reading of the manuscript, and Dr. Bingjie Zhang for assistance on scCUT&Tag-pro analysis. This work was supported by NIH grants R35GM133712 (CZ), R21HG012981 (CZ), R00CA259526 (ZS), and R01CA060499 (AD).

