## Supplementary Figures for "PATTY corrects open-chromatin bias for improved bulk and single-cell CUT&Tag profiling"

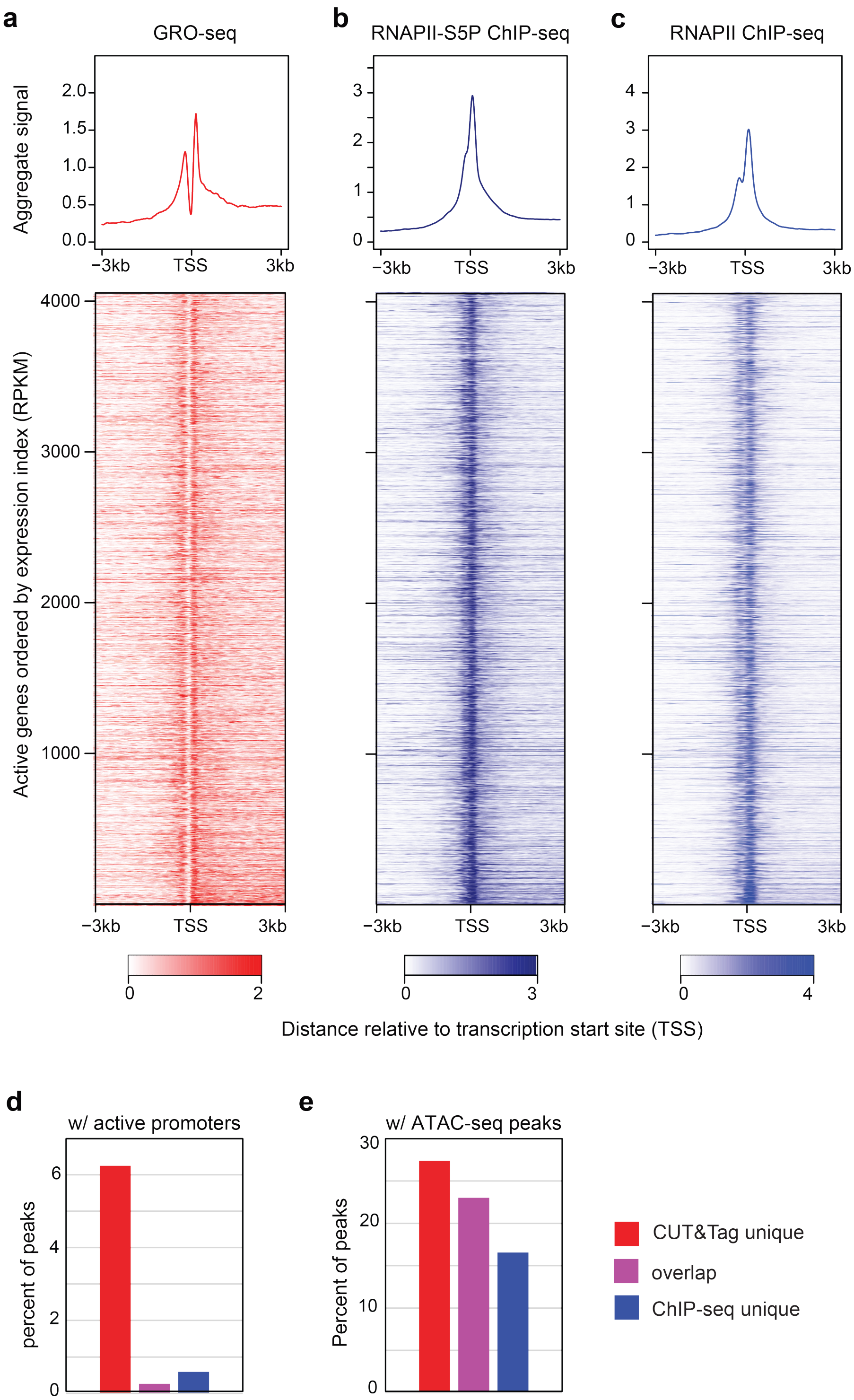


**Figure S1. Enrichment of H3K27me3 CUT&Tag signal on active gene promoters.** (**a-c**) Signal patterns across active promoter regions in K562 cells, for two GRO-seq (**a**), RNAPII-s5p ChIP-seq (**b**), and RNAPII ChIP-seq (**c**). The upper panels are the normalized aggregate signal patterns. The lower panels are heatmaps of the signal patterns at promoter regions (TSS ±3kb) of the actively transcribed genes. Rows correspond across heatmaps. (**d**, **e**) Percentage of CUT&Tag-unique (red), CUT&Tag-ChIP-overlap (purple), and ChIP-seq-unique (blue) peaks that overlap with active gene promoter regions (TSS ±1kb) (**d**) and with ATAC-seq peaks (**e**).

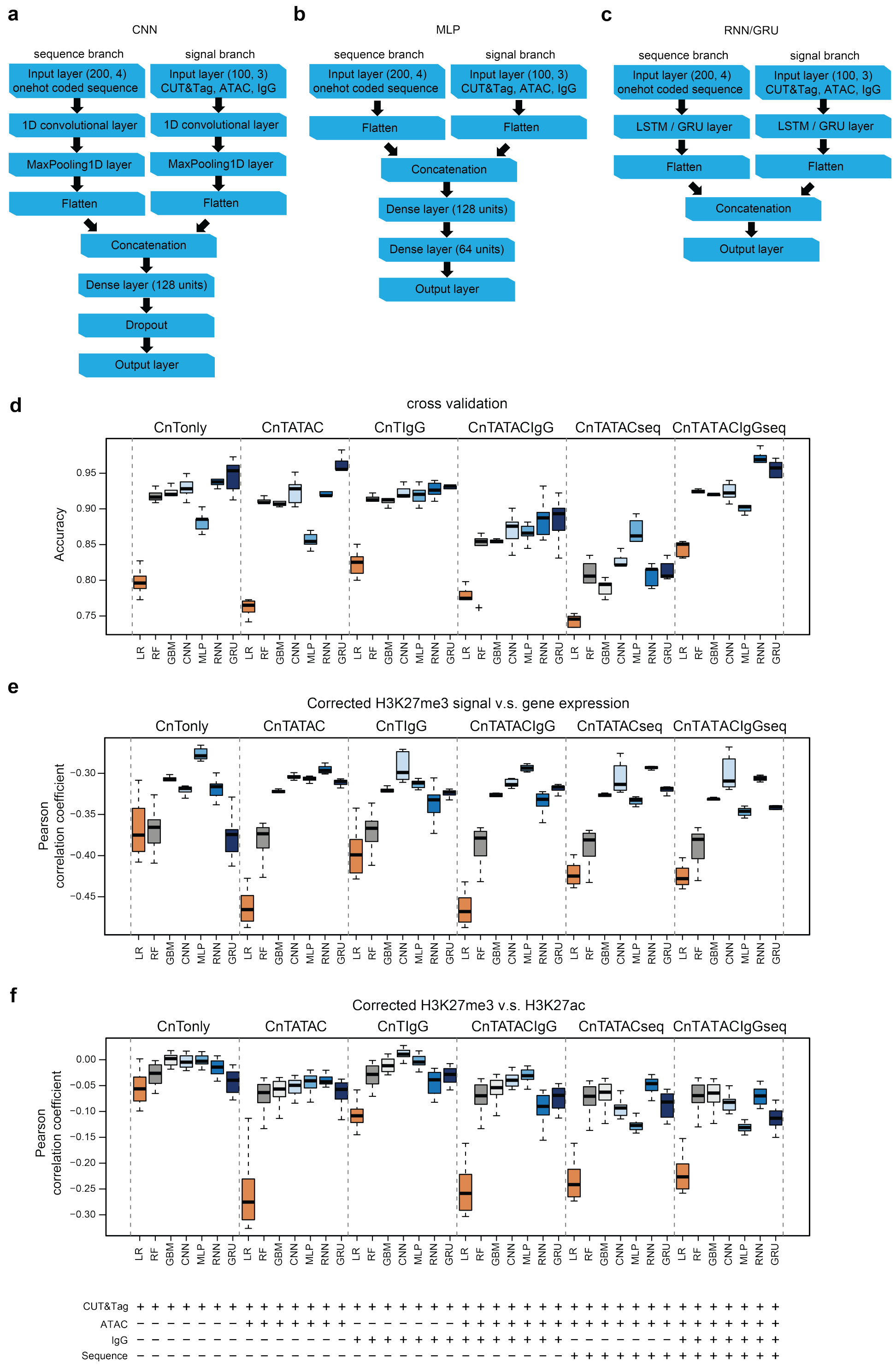


**Figure S2. Performance of different models for H3K27me3 CUT&Tag.** (**a-c**) Schematic of the (**a**) CNN, (**b**) MLP, (**c**) RNN, and GRU models. (**d**) The accuracy of different prediction models was evaluated using 5-fold cross-validation results. (**e**) Pearson correlation coefficient (y-axis) between the model-derived H3K27me3 score and gene expression across all gene promoter bins in the genome. (**f**) Pearson correlation between the model derived H3K27me3 CUT&Tag score and H3K27ac ChIP-seq signal level across all bins with CUT&Tag reads in the genome. Models with the same feature combination were displayed in the same block. Each data point in a boxplot represents a CUT&Tag sample.

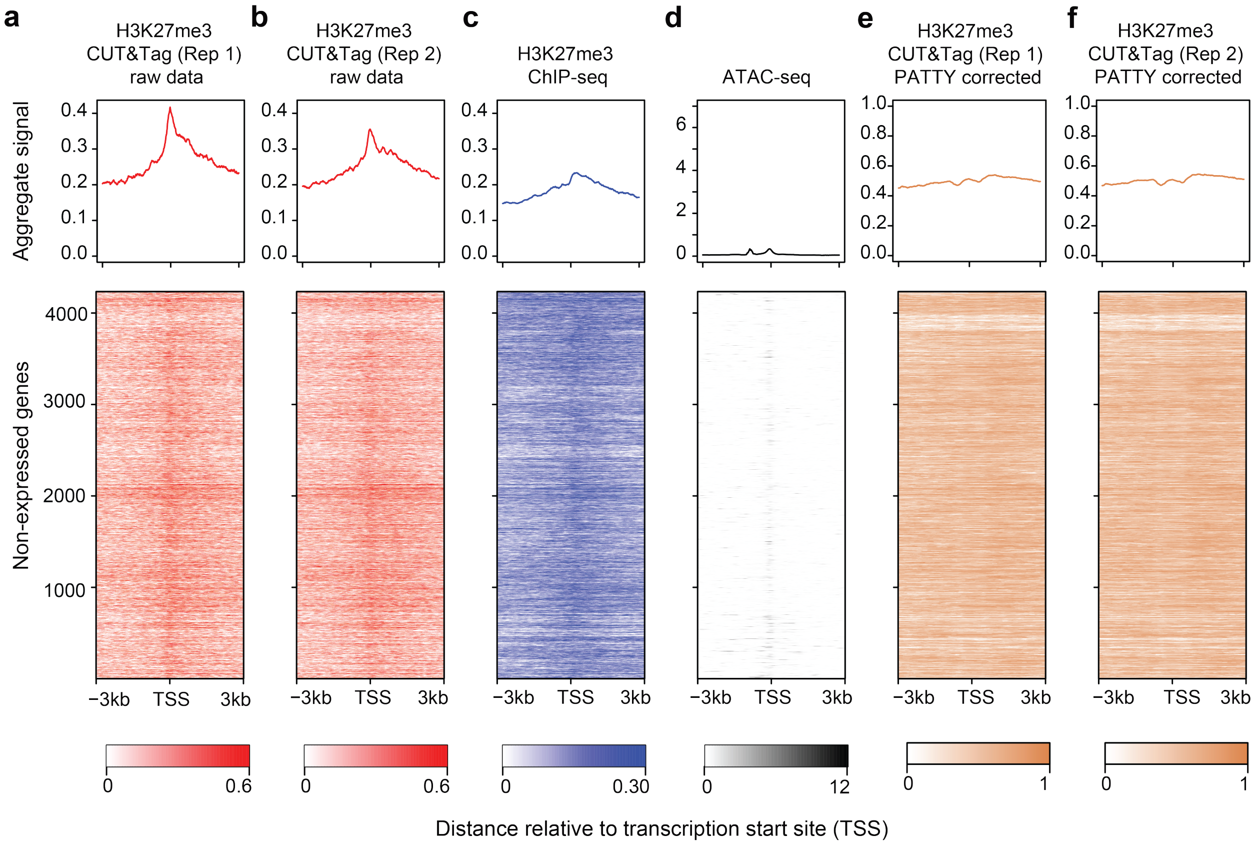


**Figure S3. H3K27me3 CUT&Tag signal patterns on repressive genes before and after PATTY correction.** (**a**-**f**) H3K27me3 CUT&Tag signal patterns across repressive gene promoter regions before (**a**, **b**) and after (**e**, **f**) bias correction by PATTY. ChIP-seq (**c**) and ATAC-seq (**d**) signals across the same regions are shown for reference. The upper panels are the normalized aggregate signal patterns. The lower panels are heatmaps of the signal patterns at promoter regions (TSS ±3kb) of the repressive genes. Rows correspond across heatmaps.

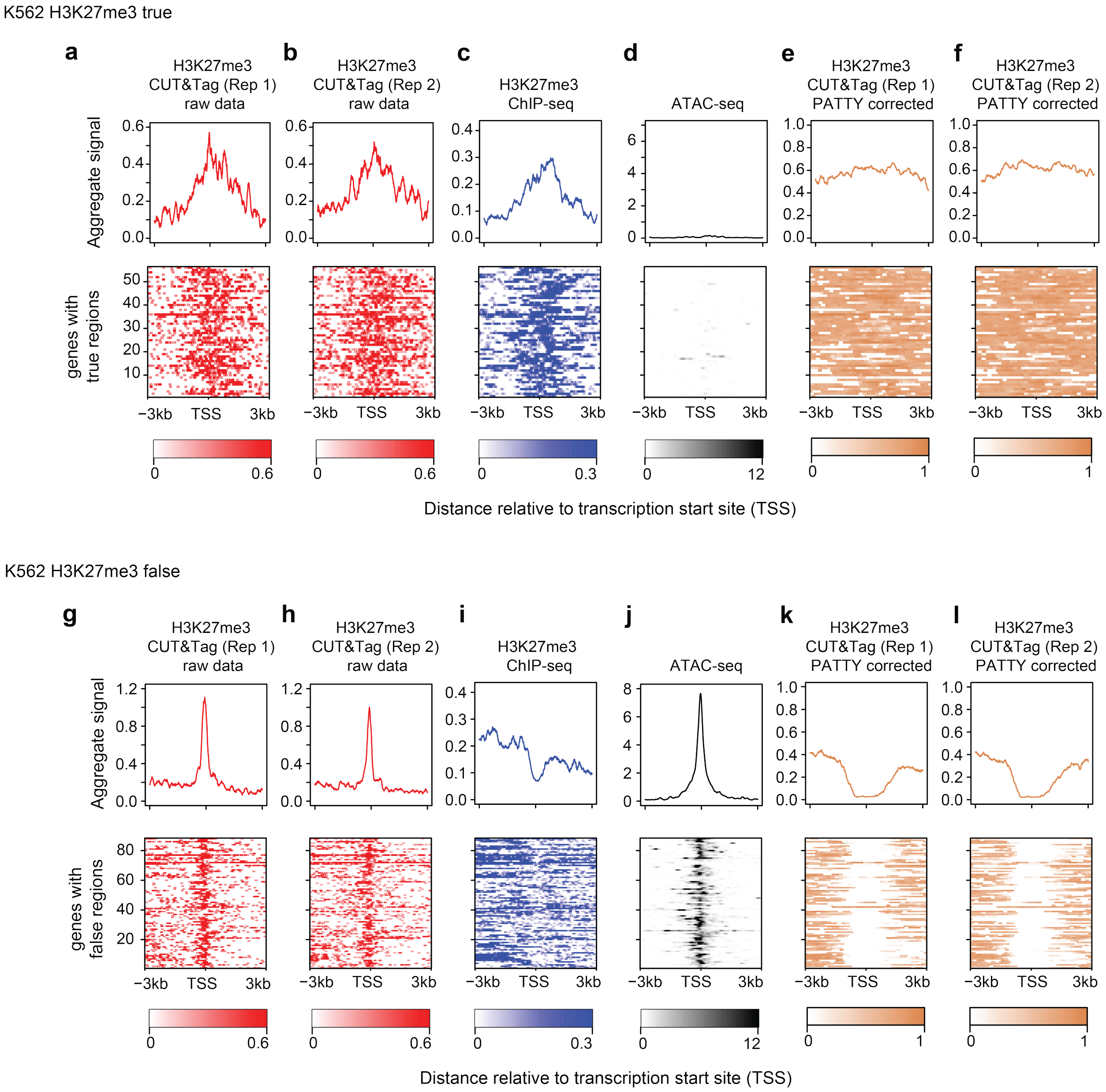


**Figure S4. PATTY correction on the H3K27me3 ground-truth regions.** Signal patterns across gene promoter regions associated with the curated true (**a**-**f**) and false (**g**-**l**) H3K27me3-marked regions. Samples include 2 replicates of H3K27me3 CUT&Tag before (**a**, **b**, **g**, **h**) and after (**e**, **f**, **k**, **l**) PATTY correction, H3K27me3 ChIP-seq (**c**, **i**), and ATAC-seq (**d**, **j**). The upper panels are normalized aggregate signal patterns. The lower panels are heatmaps of the signal patterns. Rows correspond across heatmaps.

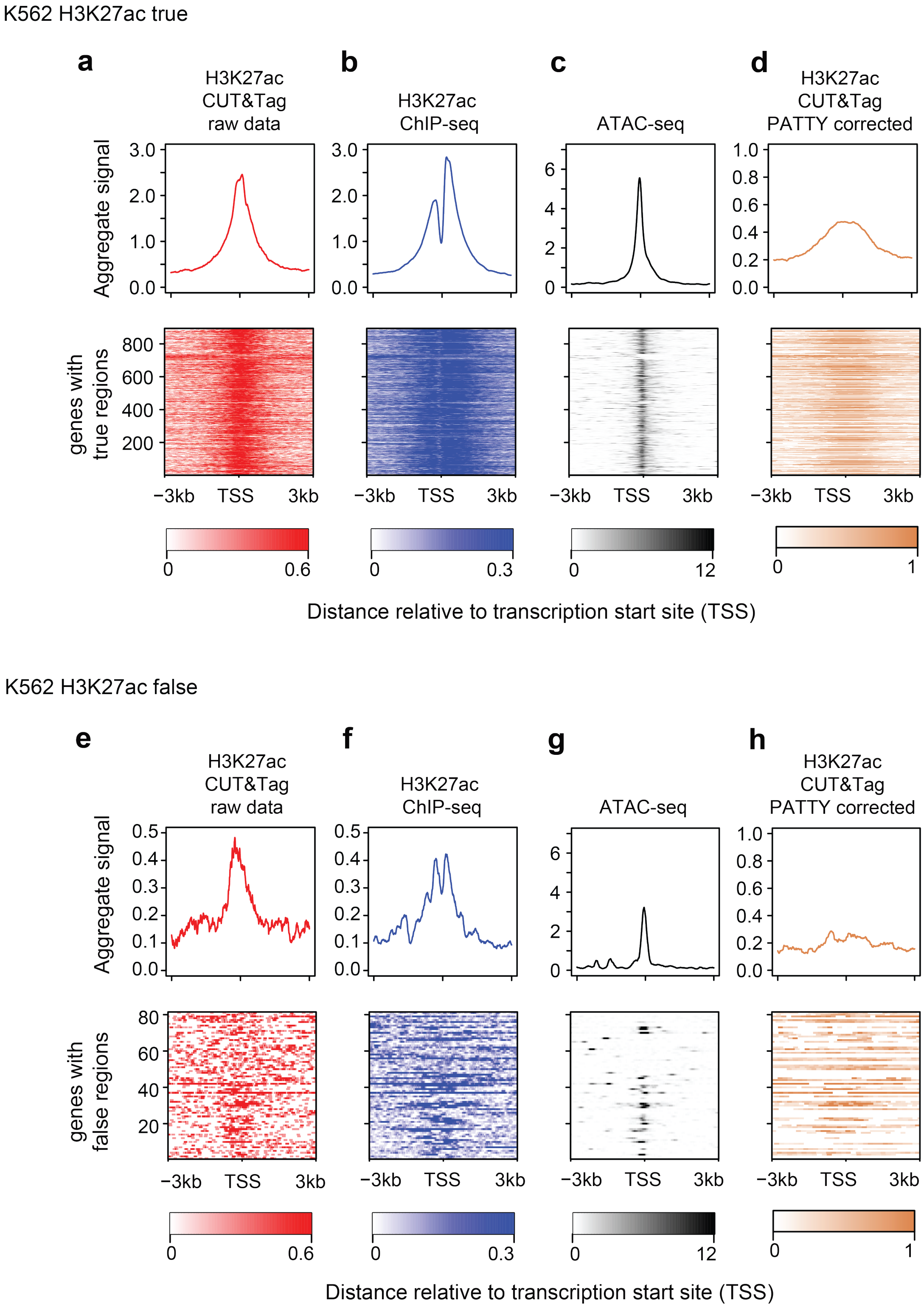


**Figure S5. PATTY correction on the H3K27ac ground-truth regions.** Signal patterns across gene promoter regions associated with the curated true (**a-d**) and false (**e-h**) H3K27ac-marked regions. Samples include H3K27ac CUT&Tag before (**a**, **e**) and after (**d**, **h**) PATTY correction, H3K27ac ChIP-seq (**b**, **f**), and ATAC-seq (**c**, **g**). The upper panels are normalized aggregate signal patterns. The lower panels are heatmaps of the signal patterns. Rows correspond across heatmaps.

K562 H3K9me3 true

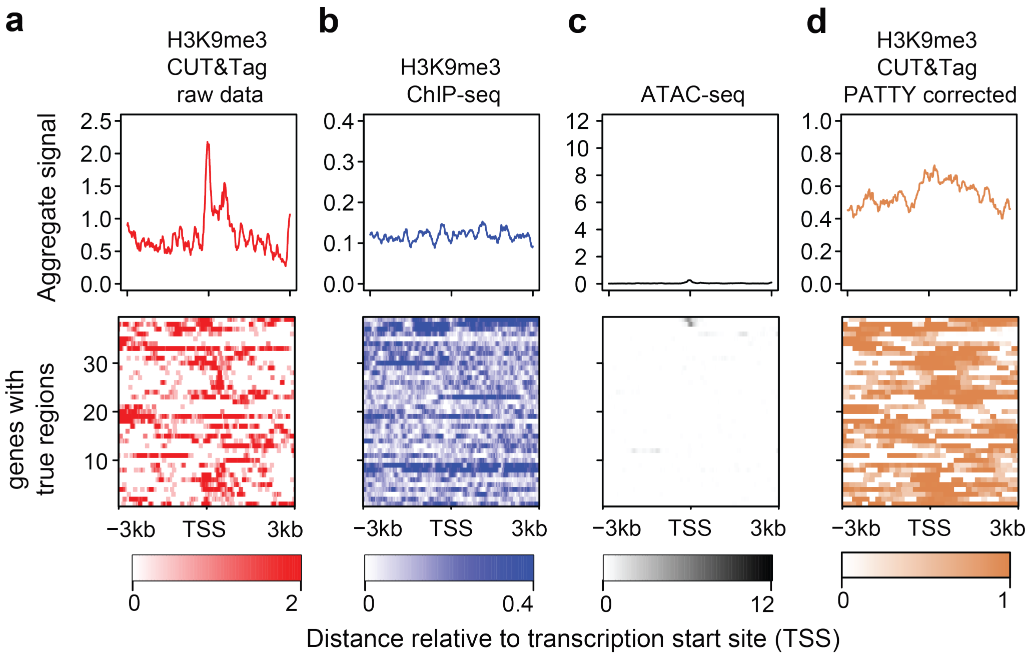


K562 H3K9me3 false

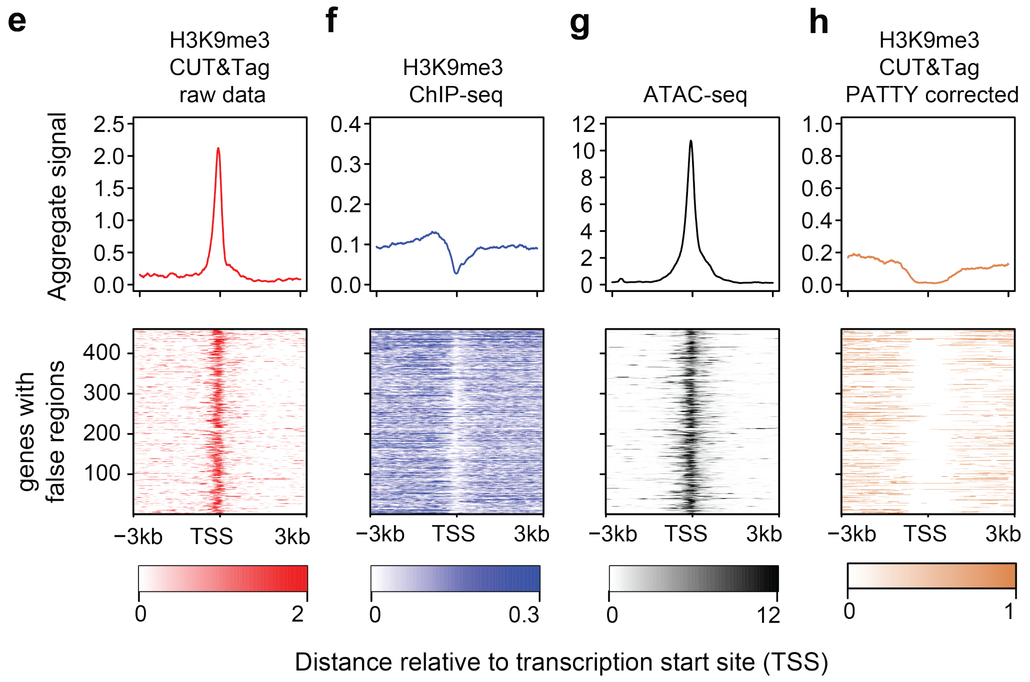


**Figure S6. PATTY correction on the H3K9me3 ground-truth regions.** Signal patterns across gene promoter regions associated with the curated true (**a-d**) and false (**e-h**) H3K9me3-marked regions. Samples include H3K9me3 CUT&Tag before (**a**, **e**) and after (**d**, **h**) PATTY correction, H3K9me3 ChIP-seq (**b**, **f**), and ATAC-seq (**c**, **g**). The upper panels are normalized aggregate signal patterns. The lower panels are heatmaps of the signal patterns. Rows correspond across heatmaps.

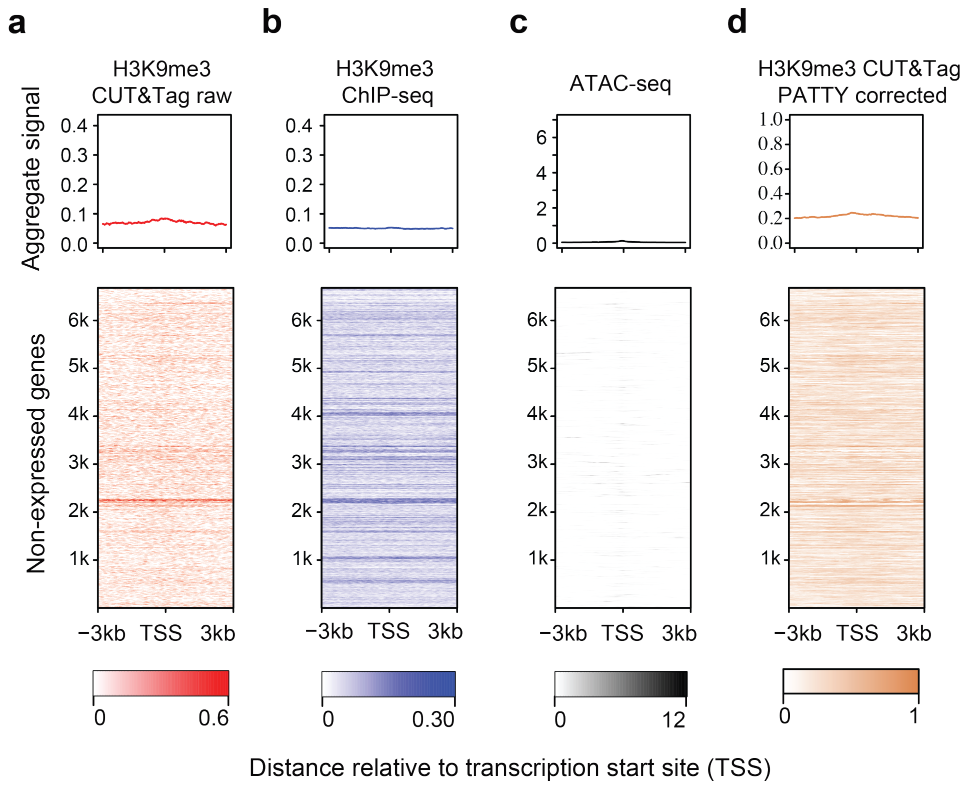


**Figure S7. H3K9me3 CUT&Tag signal patterns on repressive genes in HCT116 cell line before and after PATTY correction.** (**a-d**) H3K9me3 CUT&Tag signal patterns across repressive gene promoter regions before (**a**) and after (**d**) bias correction by PATTY. ChIP-seq (**b**) and ATAC-seq (**c**) signals across the same regions are shown for reference. The upper panels are normalized aggregate signal patterns. The lower panels are heatmaps of the signal patterns at promoter regions (TSS ±3kb) of the repressive genes. Rows correspond across heatmaps.

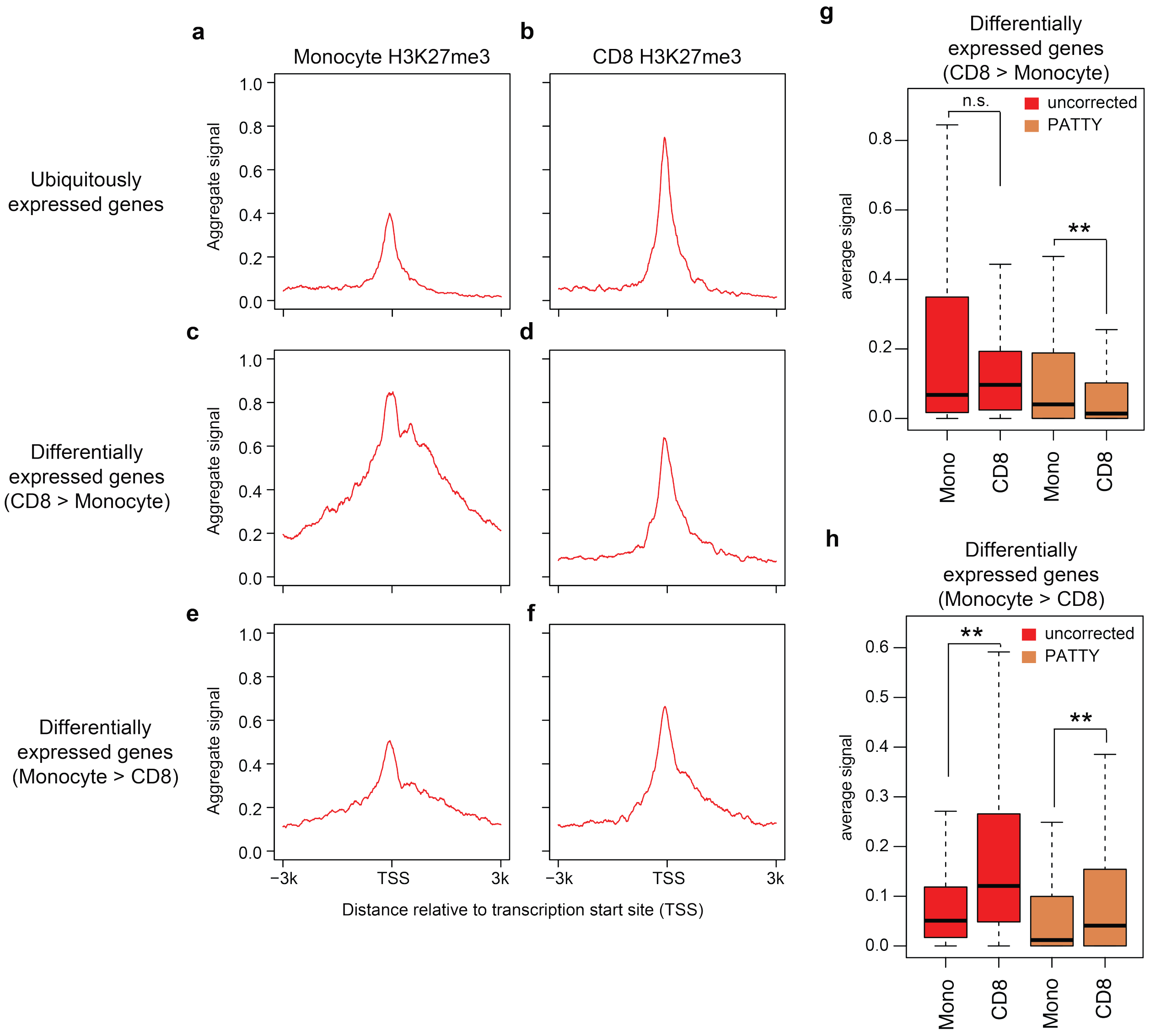


**Figure S8. H3K27me3 CUT&Tag signal in scCUT&Tag-pro data.** (**a**-**f**) Aggregate H3K27me3 scCUT&Tag-pro signal patterns across the promoter regions (TSS ±3kb) of PBMC-ubiquitously expressed genes (**a**, **b**), differentially expressed genes (DEGs) between Monocytes and CD8+ T cells that are higher in CD8 than Monocytes (**c**, **d**), and higher in Monocytes than CD8 (**e**, **f**), for scCUT&Tag-pro signal as pseudo-bulk in Monocytes (**a**, **c**, **e**) and in CD8+ T cells (**b**, **d**, **f**). (**g**, **h**) Average H3K27me3 scCUT&Tag-pro signal levels in a pseudo-bulk of cell group (labeled under x-axis) on the promoter regions (TSS ±3 kb) of DEGs high in CD8 (**g**) and DEGs high in Monocytes (**h**), comparing uncorrected data (red) with PATTY-corrected data (yellow). Each data point in a boxplot represents the H3K27me3 scCUT&Tag-pro signal for a gene. **, p < 0.01, by one-sided Wilcoxon signed-rank test.

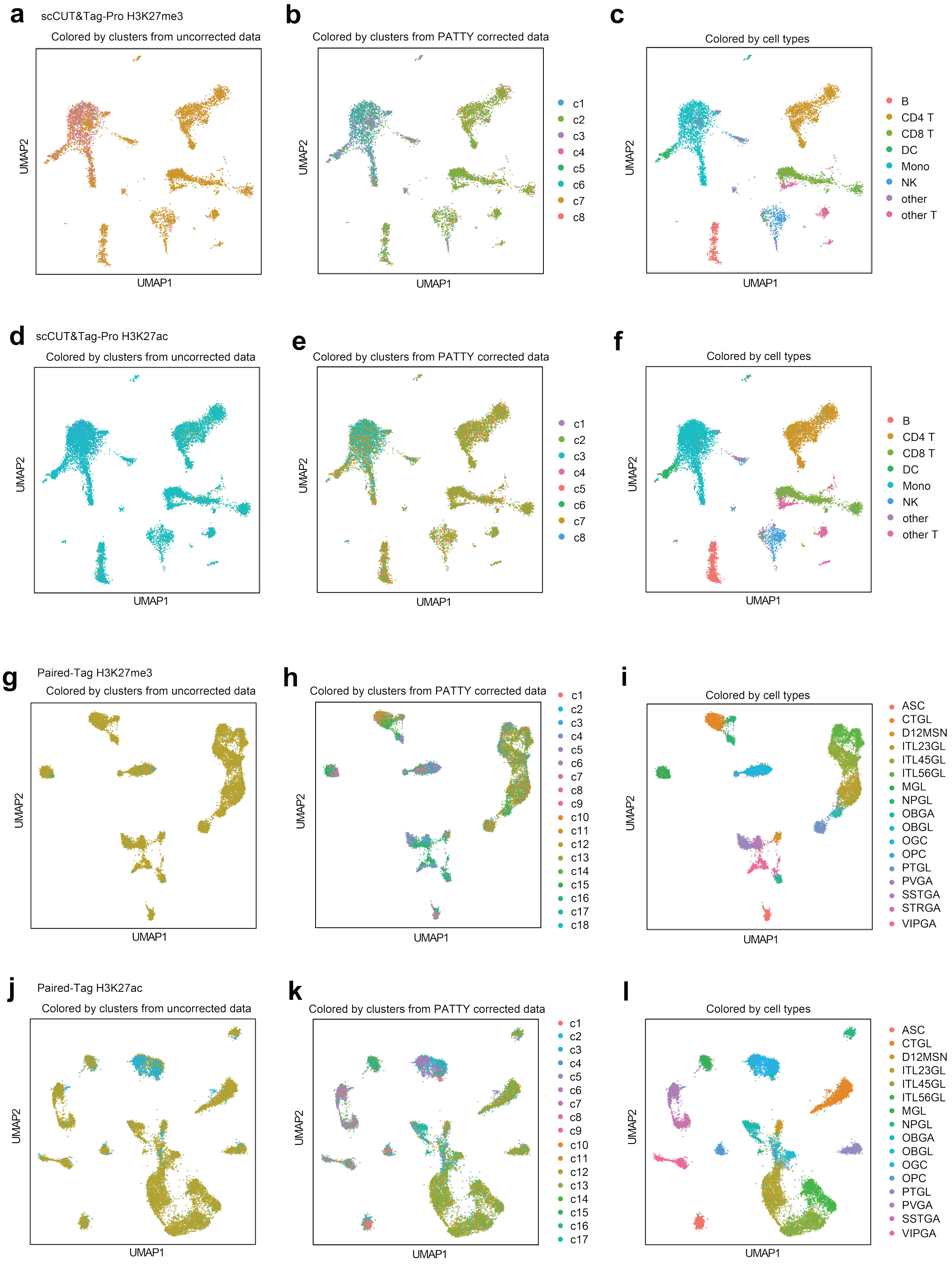


**Figure S9. Improved cell clustering on single-cell CUT&Tag after PATTY correction for scCUT&Tag-pro (a-f) and Paired-Tag (g-l) data.** (**a-l**) Visualization of H3K27me3 (**a-c**, **g-i**) and H3K27ac (**d-f**, **j-l**) single-cell using UMAP projection reported in the original publications for scCUT&Tag-pro (**a-f**) and Paired-Tag (**g-l**). Cells are colored by cluster assignments based on uncorrected data (**a**, **d**, **g**, **j**), cluster assignments based on PATTY-corrected data (**b**, **e**, **h**, **k**), or the published cell-type annotations (ground truth) (**c**, **f**, **i**, **l**).

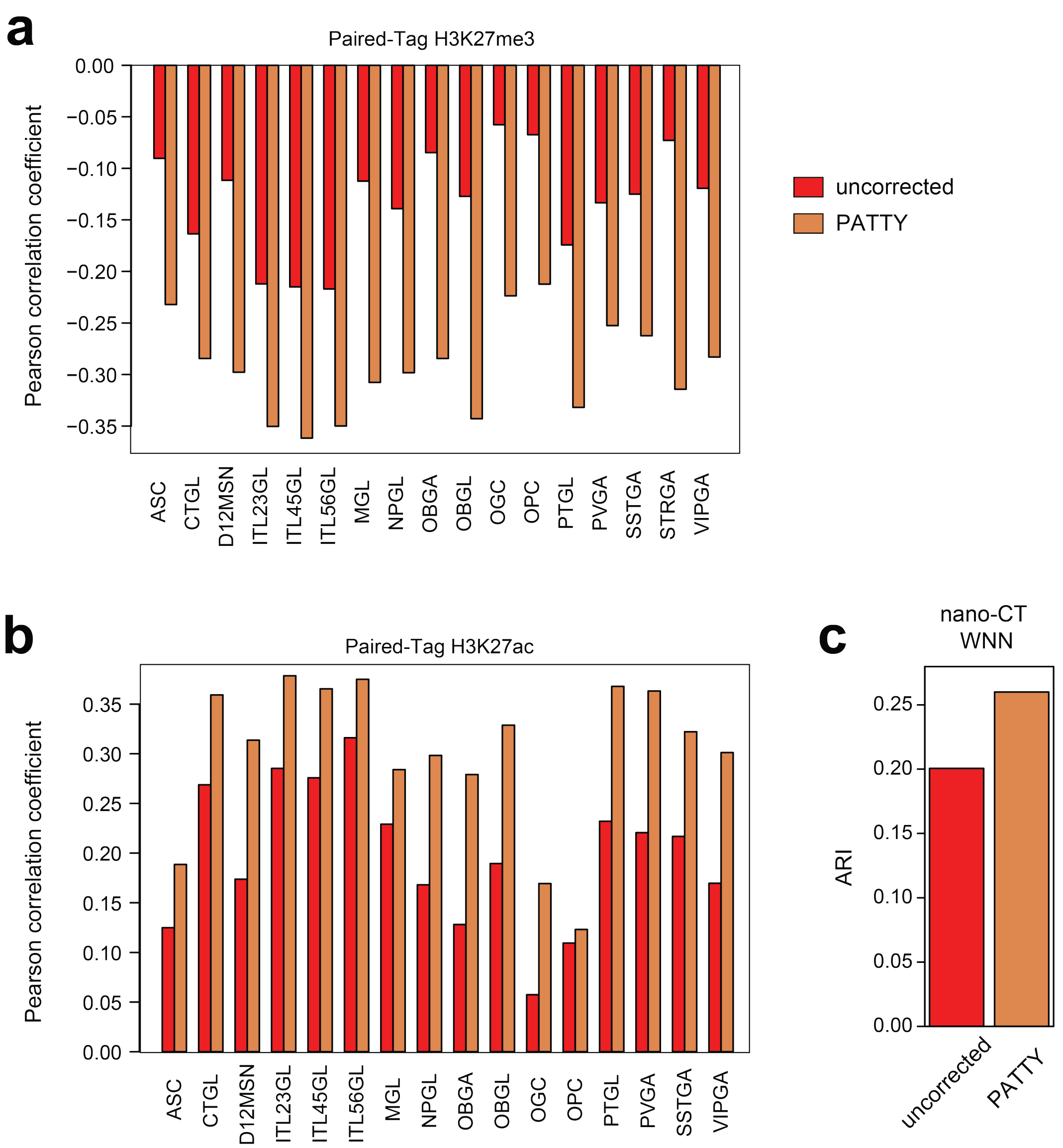


**Figure S10.** (**a**, **b**) PATTY increases concordance between histone-mark signal and gene expression for all cell types in single-cell Paired-Tag data. Bar plots showing Pearson correlation across all genes between average expression level (from RNA modality) and average H3K27me3 (**a**) or H3K27ac (**b**) signal (from CUT&Tag modality) over all cells for each cell type. Correlations using histone-mark signal from uncorrected and from PATTY-corrected data are compared side-by-side for each cell type. (**c**) Adjusted Rand index (ARI) between WNN-based clustering results and the published nano-CT cell-type annotations, comparing clustering on uncorrected and PATTY-corrected data. WNN integration was performed using H3K27me3 and H3K27ac measured in the same cells with shared barcodes.

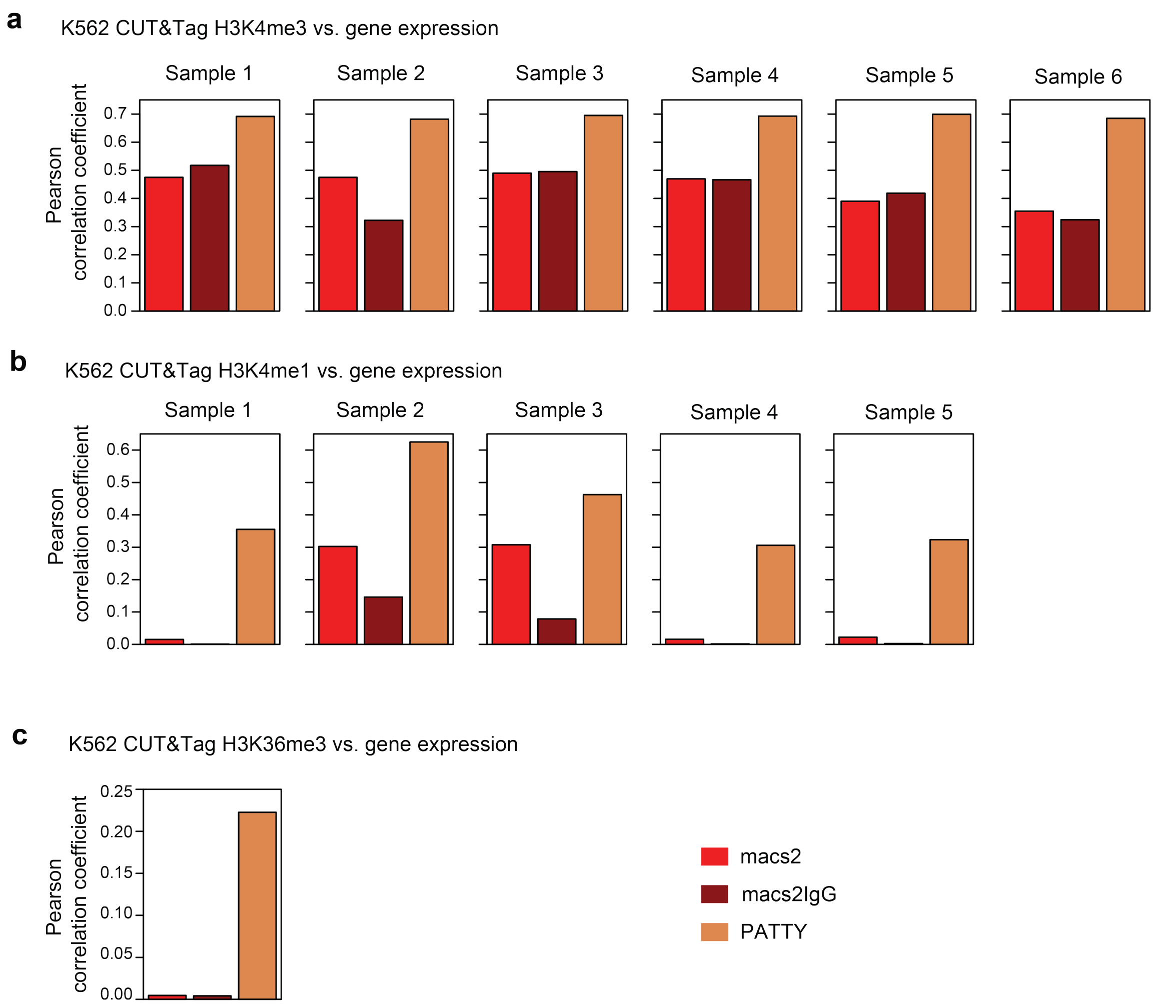


**Figure S11. Pretrained H3K27ac PATTY model improves concordance with gene expression for other active histone marks in K562.** Pearson correlation between CUT&Tag signal and gene expression across all genes in K562 for H3K4me3 (**a**), H3K4me1 (**b**), and H3K36me3 (**c**).
